# Cryo-EM Structures Reveal Tau Filaments from Down Syndrome Adopt Alzheimer’s Disease Fold

**DOI:** 10.1101/2024.04.02.587507

**Authors:** Ujjayini Ghosh, Eric Tse, Hyunjun Yang, Marie Shi, Christoffer D. Caro, Feng Wang, Gregory E. Merz, Stanley B. Prusiner, Daniel R. Southworth, Carlo Condello

## Abstract

Down syndrome (DS) is a common genetic condition caused by trisomy of chromosome 21. Among their complex clinical features, including musculoskeletal, neurological, and cardiovascular disabilities, individuals with DS have an increased risk of developing progressive dementia and early-onset Alzheimer’s disease (AD). This dementia is attributed to the increased gene dosage of the amyloid-β (Aβ) precursor protein gene, the formation of self-propagating Aβ and tau prion conformers, and the deposition of neurotoxic Aβ plaques and tau neurofibrillary tangles. Tau amyloid fibrils have previously been established to adopt many distinct conformations across different neurodegenerative conditions. Here, we report the characterization of brain samples from four DS cases spanning 36 to 63 years of age by spectral confocal imaging with conformation-specific dyes and cryo–electron microscopy (cryo-EM) to determine structures of isolated tau fibrils. High-resolution structures revealed paired helical filament (PHF) and straight filament (SF) conformations of tau that were identical to those determined from AD cases. The PHFs and SFs are made of two *C*-shaped protofilaments, each containing a cross-β/β-helix motif. Similar to filaments from AD cases, most filaments from the DS cases adopted the PHF form, while a minority (approximately 20%) formed SFs. Samples from the youngest individual with no documented dementia had sparse tau deposits. To isolate tau for cryo-EM from this challenging sample we used a novel affinity-grid method involving a graphene oxide surface derivatized with anti-tau antibodies. This method improved isolation and revealed that primarily tau PHFs and a minor population of chronic traumatic encephalopathy type II–like filaments were present in this youngest case. These findings expand the similarities between AD and DS to the molecular level, providing insight into their related pathologies and the potential for targeting common tau filament folds by small-molecule therapeutics and diagnostics.

## INTRODUCTION

Down syndrome (DS) is the most common chromosomal disorder of intellectual disability, affecting approximately 400,000 people in the United States and approximately 5.5 million people worldwide[1]. DS results from the presence of a third copy of chromosome 21 (Chr21) and associated genetic dosage changes that cause varying and complex disabilities, including heart defects, obesity, and diabetes, throughout development and adulthood. With recent increases in the lifespan of persons with DS, it is now realized that Alzheimer’s disease (AD) is a substantial comorbidity in individuals with DS that arises at a much earlier age than other populations[2]. Nearly all adults with DS develop severe AD neuropathological changes by age 40; however, there is evidence that the first appearance of plaques and tangles occurs as early as the teens or twenties[3, 4]. Further investigation is needed in order to understand the molecular pathology of early-onset AD resulting from increased amyloid-β (Aβ) precursor protein (APP) production in DS.

As with familial and sporadic forms of AD, extracellular Aβ plaques and intraneuronal tau neurofibrillary tangles (NFTs) are hallmarks of AD that arise in DS and thus are strongly implicated in neurodegeneration and cognitive decline. Early-onset Aβ plaque deposition occurs from increased expression of the *APP* gene and resulting overabundance of its cleavage product, Aβ42, given APP is present on Chr21. Increases in hyperphosphorylated tau and NFTs are also likely linked to increased Aβ42, considering gene duplication of *APP* is sufficient to cause early-onset AD[5]. Conversely, a small fraction of DS cases have only a partial trisomy of Chr21 and lack the extra copy of the *APP* gene; these cases do not develop the neuropathological changes of AD[6, 7]. Taken together, these findings bolster the notion that APP overexpression is the major catalyst for AD neuropathology in older individuals with DS. While the clinical and biomarker similarities are now well established between sporadic AD (sAD) that arises in general populations and AD in individuals with DS[8–10], the molecular pathology is less well understood, given the substantial gene dosage changes with trisomy 21. Previously, we have shown that self-propagating prion conformers of Aβ and tau accumulate in abundance in DS brains with increasing age[11]; in contrast, sAD samples showed the opposite trend with a decreasing abundance of Aβ and tau prions in long-lived patients despite an overall increase in total insoluble tau[12]. Thus, although there is strong evidence for the overall similarity in the histopathological and clinical biomarkers of tau NFT accumulation between DS and AD, it is not known if the molecular structures of the associated filaments are similar.

Recent breakthroughs using cryo–electron microscopy (cryo-EM) have revealed striking differences between tau filament structures isolated from cases with sAD, Pick’s disease, corticobasal degeneration, progressive supranuclear palsy, and several other primary and secondary tauopathies[13–18]. Alternative splicing of tau results in isoforms that differ in the number of N-terminal regions and microtubule-binding repeat regions and thus exhibit distinct interactions of tau β sheets, resulting in different folds in each of these diseases. Both the three-repeat and four-repeat tau isoforms implicated in AD form paired helical filaments (PHFs) in AD cases, whereas Pick’s disease presents with the predominant amyloid structures as single filaments, and progressive supranuclear palsy presents as enriched in three-repeat and four-repeat tau isoforms. Corticobasal degeneration is also a four-repeat tauopathy and presents both single and paired filaments. These findings indicate that tau filament conformations are disease specific. Thus, propagation of a particular tau filament fold is likely linked to specific cellular and molecular factors associated with different disease pathologies[18]. We postulate that overexpression of the 250 protein-coding genes on Chr21 in DS and the high prevalence of early-onset AD present unique cellular drivers of tau filament propagation. Additionally, given the importance of developing structure-guided molecular diagnostics and therapeutics that target tau filaments to more precisely define pathologies that arise in DS and in neurodegenerative diseases, we were motivated to characterize tau filament structures that develop in individuals with DS.

Here we sought to characterize and determine cryo-EM structures of tau filaments isolated from several postmortem tissue samples from individuals with DS across different ages. Tau filaments were successfully purified from four individuals, aged 63, 51, 46, and 36 years (cases 1 to 4, respectively). Cases 1, 2, and 3 showed robust tau pathology similar to end-stage sAD. However, samples from case 4, the 36-year-old, exhibited sparse tau deposits. Tau PHF and straight filament (SF) structures from the 63-year-old case were determined to 2.7-Å and 2.9-Å resolution, respectively, establishing complementarity to AD tau filament conformations at high resolution. Moreover, structures from all four individuals demonstrated that tau adopts the canonical PHF and SF forms, and these forms are present at similar percentages to what has been observed for AD. To overcome the low level of tau filaments obtained from the 36-year-old case, we used a custom-developed graphene oxide (GO) affinity-grid method in which the cryo-EM sample grids were derivatized with anti-tau antibodies to capture and enrich for tau filaments prior to vitrification. Thus, we present an “on-grid” tau filament isolation method that may be broadly useful for structure determination of filaments from low-abundance sources. Overall, these results establish that, despite substantial differences in genetic background and age, tau filaments that arise in individuals with DS and neurodegeneration are identical to those in AD. Our data provide the molecular framework required to identify conformation-specific diagnostic probes and to design novel therapeutic compounds targeting tau polymorphs found in DS.

## METHODS

### Clinical history and neuropathology

Deidentified human biospecimens from deceased individuals were obtained from academic biorepositories. All tissue donors provided written or verbal consent to donate autopsied brains for use in biomedical research in accordance with the standards of each institution. This study was exempt from institutional review board approval (i.e., this study is not considered human subject research) in accordance with the University of California San Francisco institutional review board policy. We determined the cryo-EM structure of tau filaments from 0.5 g fresh-frozen frontal cortex samples from four individuals with DS. Case 1 was a 63-year-old man with documented dementia. Case 2 was a 51-year-old woman with documented dementia. Case 3 was a 46-year-old woman with no documented dementia. Case 4 was a 36-year-old woman with no documented dementia. The demographic and cliniconeuropathological details of these four cases are in **Supplementary Table S1**. Formalin-fixed samples from additional cases of DS and sAD were obtained for histological studies. Fresh-frozen samples from additional cases of DS and sAD were obtained for biochemical studies. The demographic and cliniconeuropathological details of the additional cases are in **Supplementary Table S2** and **Supplementary Table S3**.

### Immunohistochemistry

Deparaffinized fixed sections were pretreated in 98% formic acid for 6 minutes to enhance immunoreactivity. After blocking with 10% normal goat serum in phosphate-buffered saline (PBS) with 0.2% Tween 20 (PBST), sections were incubated at room temperature in primary antibodies overnight followed by secondary antibodies for 2 hours. Primary antibodies were prepared in 10% normal goat serum and applied as combinations of either (a) anti-Aβ1-40 rabbit polyclonal (1:200; AB5074P, MilliporeSigma) and anti-Aβ1-42 12F4 mouse monoclonal (1:200; 805502, BioLegend); (b) anti-Aβ17-24 4G8 mouse monoclonal (1:1000; 800709, BioLegend) and anti-tau (phospho-S262) rabbit polyclonal (1:200; ab131354, Abcam); or (c) anti-tau AD conformation-specific GT-38 mouse monoclonal (1:150; ab246808, Abcam) and anti-tau (phospho-S396) rabbit monoclonal (1:200; ab109390, Abcam) antibodies. The polyclonal IgG H+L secondary antibodies were conjugated with Alexa Fluor 488 or 647 (A11029, A21235, A11008, and A21244, Thermo Fisher Scientific) applied 1:500 in 10% normal goat serum in PBST. Stained slides were scanned on a ZEISS Axio Scan Z1 digital slide scanner at 20× magnification. Excitation at 493, 553, or 653 nm was followed by detection at 517 nm (Aβ40 or phospho-tau), 568 nm (autofluorescence), or 668 nm (Aβ42 or tau), respectively.

### Dye staining of brain sections for EMBER (*Excitation Multiplexed Bright Emission Recordings*) microscopy

Formalin-fixed, paraffin-embedded brains were sectioned (8-μm thickness) and glass mounted. To reduce the autofluorescence in the brain tissue by greater than 90% intensity (e.g., from lipofuscin or hemosiderin), the sections were photobleached in a cold room for up to 48 hours using a multispectral LED array[19]. The sections were deparaffinized, washed in PBS, and stained with 5 µM dye 60 for 30 minutes. The sections were washed with PBS and coverslipped with PermaFluor (Thermo Scientific) as the self-curing mounting medium. See Yang et al. for additional details[20].

### EMBER microscopy data collection

Dye 60-stained fixed brain slices were imaged with a Leica Microsystems SP8 confocal microscope using a 40× water-immersion lens (1.1 NA), a white light and 405 nm lasers, and a HyD detector at 512 × 512-pixel resolution at 2× zoom. The optical plane was manually focused for each field of view. To reduce the background noise from the bottom of the slide, the LightGate was set to 0.5 to 18 ns. First, 110 images were acquired using the Λ/λ-scan mode with wavelength excitations of 470, 490, 510, 530, 550, 570, 590, 610, 630, 650, and 670 nm. The emission detection range started at 10 nm greater than the given excitation wavelength and ended at 780 nm, with a 20-nm window. For example, for the 470-nm excitation, the images were collected at 480 to 500, 500 to 520, 520 to 540, 540 to 560, 560 to 580, 580 to 600, 600 to 620, 620 to 640, 640 to 660, 660 to 680, 680 to 700, 700 to 720, 720 to 740, 740 to 760, and 760 to 780 nm. Then, in the λ-scan mode, 18 additional images were collected at 405-nm excitation with emission detection intervals of 20 nm from 420 nm to 780 nm. See Yang et al. for additional details[20].

### EMBER data postprocessing, particle segmentation, and dimension reduction

The EMBER analytical pipeline is described in detail in Yang et al. [20]. In brief, we developed a set of custom scripts in MATLAB to process the raw fluorescent images and segment the aggregated protein deposits. The signal-processing algorithm for the analysis of particle-resolution EMBER spectra was executed in MATLAB with *pca*. Each identified EMBER particle from the particle segmentation was normalized to [0, 1] and then concatenated in an array for principal component analysis. The principal component scores PC1 and PC2 were plotted. Uniform manifold approximation and projection (UMAP) was performed in MATLAB with *run_umap* with default settings.

### Preparation of sarkosyl-insoluble tau for biochemical characterization

Sarkosyl-insoluble Tau was prepared by diluting 250uL 10% brain homogenate in equal volume of A68 buffer (10 mM Tris-HCl, pH 7.4, 0.8 M NaCl, 1 mM EGTA, 5 mM EDTA, 10% sucrose, protease and phosphatase inhibitors), followed by centrifugation at 13,000 × g at 4 °C for 20 min^1^. The supernatants were kept on ice, and the pellets were resuspended in 250 μl of A68 buffer and centrifuged for another 20 min. Both supernatants were pooled. Samples were then incubated with 1% sodium lauroyl sarcosinate (1614374, Sigma Aldrich) at room temperature for 1 h at 700 rpm followed by centrifugation at 100,000 × g at 4 °C for 1 h. Pellets were then washed and resuspended in 50 mM Tris-HCl, pH 7.4 at half the original volume of brain homogenate. Samples were used immediately or snap-frozen in liquid nitrogen and stored at −80 °C.

### Homogenous time-resolved fluorescence assay (HTRF) for tau aggregation

HTRF was performed on sarkosyl-insoluble extracts using Tau Aggregation Kits (6FTAUPEG, Revvity) with 384-well microplates (Perkin Elmer) per manufacturer instructions. Briefly, anti-human TAU-d2 conjugate and anti-human TAU-Tb^3+^-cryptate conjugate antibody stocks were diluted at a concentration of 1:19 with detection buffer and premixed 1:1 immediately before plating. Samples were prepared by diluting sarkosyl-insoluble extracts at a concentration of 1:4 with 50mM tris buffer. Samples (10uL) were then added to each well followed by 10uL of premixed antibodies. Plates were sealed and incubated for 2 h at room temperature. Plates were read at 665 nm and 620 nm using the PHERAstar FSX Microplate Reader followed by analysis using MARS Data Analysis Software.

### Western blot analysis

Samples were selected based upon normalized insoluble tau protein content as determined by the HTRF Tau Aggregation assay. Sarkosyl-insoluble tau samples were treated with or without trypsin (final trypsin concentration 2mg/mL) at room temperature for 1 h at 1400 rpm. The samples were then resolved using 4–15% precast polyacrylamide gels and then transferred to a nitrocellulose membrane. Briefly, samples were mixed with 4X Laemlli sample buffer (1610747, Bio-Rad) and 2-mercaptoethanol followed by boiling for 5 minutes for protein denaturation. Following membrane transfer and blocking, tau protein was probed using the following primary antibody pairs: 1) T13 tau (1:1000, sc-21796, Santa Cruz) and 4R tau (1:1000, ab218314, Abcam); 2) GT-38 (1:1000, ab246808, Abcam) and TauC4 (1:1000, ABN2178, Sigma-Aldrich). Membranes were incubated with primary antibody pairs overnight at 4 °C with rocking; all primary antibody cocktails were prepared in Intercept blocking buffer (LI-COR). Following a TBST wash, membranes were incubated with goat anti-rabbit secondary antibody conjugated with IRDye 680RD (1:10,000, #926-68071, LI-COR) and goat anti-mouse secondary antibody conjugated with IRDye 800CW (1:10,000, #926-32210, LI-COR) in Intercept blocking buffer for 1 h at room temperature. Fluorescence was detected with the Odyssey Fc Imager (LI-COR) using channels 700 and 800. The PageRuler Plus pre-stained protein ladder (PI26619, Thermo Scientific) was used to provide a molecular weight marker. Images were analyzed with the LI-COR Acquisition Software.

### Filament purification

For case 1, tau filaments were purified from the frontal cortex as previously described in Fitzpatrick et al.[13]. For cases 2 to 4, we achieved improvements in the tau filament purification, including reduced background ferritin contamination, by following methods previously described for purification of Aβ filaments[21]. Here, 0.5 g brain tissue was homogenized in 20 volumes (w/v) of extraction buffer and brought to 2% sarkosyl for incubation at room temperature for 1 hour. The homogenates were then centrifuged at 10,000 × *g* for 10 minutes, and the resulting supernatants were additionally spun at 100,000 × *g* for 60 minutes at 4°C. The pellet from the second spin was resuspended in 1 mL/g extraction buffer and centrifuged at 3000 × *g* for 5 minutes. The supernatants were diluted three-fold in buffer containing 50 mM Tris-HCl, pH 7.4, 150 mM NaCl, 10% sucrose, and 0.2% sarkosyl and centrifuged at 100,000 × *g* for 30 minutes. The resulting sarkosyl-insoluble pellets were resuspended in 100 µL/g 20 mM Tris-HCl, pH 7.4, containing 50 mM NaCl, prior to vitrification for cryo-EM.

### Affinity-grid preparation, vitrification, and data collection

GO deposition and assembly of the affinity grids was performed essentially as described[22, 23]. Briefly, to functionalize the GO surface, grids (Au QUANTIFOIL, R 1.2/1.3, 300 mesh, Quantifoil Micro Tools GmbH) were submerged in dimethyl sulfoxide with 10 mM dibenzocyclooctyne-PEG-amine (DBCO-PEG4-amine; CCT-A103P, Vector Laboratories) overnight at room temperature, washed, and then incubated with 20 µL of a 1 mM solution of azide polyethylene glycol (PEG) maleimide and methoxyl PEG azide (5 kDa and 2 kDa molecular weight, respectively; Nanocs) at a 1:9 ratio for 6 hours. Maleimide-functionalized grids were then washed with water and then ethanol, dried, and stored at −20°C. Prior to vitrification, GO grids were first incubated with 3 µL 250 nM recombinant protein G (Z02007, GenScript) in resuspension buffer (100 mM KCl and 40 mM HEPES, pH 7.4) for 15 minutes in a Vitrobot Mark IV System (Thermo Fisher Scientific) at 100% relative humidity and then washed and inactivated with 20 mM 2-mercaptoethanol in the resuspension buffer. Then, 3 µL anti-phospho-tau AT8 (S202 and T205) monoclonal antibody (1:100; MN1020, Invitrogen) was applied for 15 minutes, and then the grids were washed three times. Next, 3 µL purified tau filaments from case 2 or 4 were applied, and the grids were incubated for 15 minutes; then the grids were washed and mechanically blotted for 3 to 4 seconds and plunge frozen. For cases 1 and 3, conventional methods were used: 3 µL purified tau filaments were applied to glow-discharged holey carbon grids and mechanically blotted for 3 to 4 seconds at 100% relative humidity and plunge frozen. For all datasets, super-resolution movies were collected using a Titan Krios microscope (Thermo Fisher Scientific) operating at 300 kV and equipped with a K3 direct electron-detection camera (Gatan, Inc.) with a BioQuantum energy filter (Gatan, Inc.) with a slit width of 20 eV. Super-resolution movies were recorded at 105,000× magnification (pixel size: 0.417 Å/pixel) with a defocus range of 0.8 to 1.8 µm and a total exposure time of 2.024 seconds fractionated into 0.025-second subframes.

### Helical reconstruction

All data-processing steps were done in RELION[24]. The movies were motion-corrected using MotionCor2[25] and Fourier-cropped by a factor of two to give the final pixel size of 0.834 Å per pixel. The dose-weighted summed micrographs were directly imported to RELION[26, 27] and were used for further processing. Contrast transfer function (CTF) was estimated using CTFFIND 4.1[28]. The filaments were picked manually with a box size of 1120 pixels downsampled to 280 pixels. 2D classification was used to separate homogeneous segments for further image processing. After the first round of 2D classification, image alignment was performed to correct for the variable in-plane rotation of the fibril projections, as previously described[29]. For each well-resolved 2D class, the angle α between the *x* axis of the image and fibril-growth direction and the displacement δ of the fibril along the *y* axis from the center of the box were measured in ImageJ[30]. The nonzero values for α and δ were corrected using two MATLAB codes, as follows:

AnglePsi = AnglePsi + α
AnglePsiPrior → AnglePsiPrior + α
OriginX → OriginX + δ sin(AnglePsi)
OriginY → OriginY + δ cos(AnglePsi)

The new values of these parameters were updated in the “particles.star” file. Following the manual alignment, further 2D classification was performed to make sure all the 2D classes were well centered and aligned in the boxes. 3D initial models were then created de novo from the 2D class averages using an estimated helical rise of 4.75 Å and the crossover distances of the filaments estimated from the 2D classes with 1120-pixel boxes using RELION’s *relion_helix_inimodel2d* feature. The 3D initial models were low-pass filtered to 10 Å for further 3D classification. The particles corresponding to each initial model were re-extracted with 280-pixel boxes for further 3D classification. The best particles were selected from the 3D classification for each dataset. The final rise and twist were optimized in 3D autorefinement. The final 3D densities were sharpened using postprocessing in RELION, and the final resolution was determined from Fourier shell correlation at 0.143 Å from two independently refined half-maps using a soft-edged solvent mask. The final resolutions of tau PHF were 2.7 Å for case 1, 3.1 Å for case 2, 2.9 Å for case 3, and 5 Å for case 4. The final resolutions of tau SF were 3.0 Å for case 1, 3.2 Å for case 2, and 3.1 Å for case 3. For case 4, the resolution of the minor population of chronic traumatic encephalopathy (CTE)–like filaments was 7.8 Å. Imaging and reconstruction statistics are summarized in **Supplementary Table S4**.

### Model building and refinement

For PHF and SF model building and refinement, previous models[31] were initially docked into the final maps for cases 1 and 2. A single strand of previously solved PHF (Protein Data Bank ID: 7NRQ) or SF (Protein Data Bank ID: 7NRS) was used to rigid-body fit with the final postprocessed density map in ChimeraX[32]. An initial round of real-space refinement was performed using PHENIX[33], after which the model was used against a single half-map in ChimeraX with the ISOLDE plugin[34] to check for clashes and rotamers. A final round of PHENIX real-space refinement was performed, and MolProbity[35] statistics for each model are summarized in **Supplementary Table S5**.

## RESULTS

### Fluorescence characterization of DS brain samples from individuals aged 36 to 63 years

In prior work, we extensively characterized a large cohort of postmortem DS brain samples from several biorepositories in the United States and Europe using the following assays: (a) biochemical measurements of Aβ and tau species in soluble and insoluble fractions of bulk tissue homogenate, (b) immunohistochemical measurements of Aβ and tau species, and (c) cell-based prion bioassays for self-propagating Aβ and tau species[11, 36]. In addition to the demographic and cliniconeuropathological details provided by each biorepository, we used our rich phenotypic dataset to prioritize a list of candidate samples from the medial frontal cortex for tau filament purification and structural characterization (see **Supplementary Table S1**).

Here, our goal was to isolate and structurally characterize tau filaments from the frontal cortex of individuals with DS who died at different ages to span the neuropathological stages of AD that can develop in DS. We present the histological evaluation of samples that led to sufficient tau filament purification and structure determination by cryo-EM. Using fixed samples adjacent to the frozen samples used for purification in a given DS case, we performed a series of immunofluorescent stains for Aβ and tau species.

In DS cases 1 (63 y), 2 (51 y), and 3 (46 y), we observed abundant and widespread 4G8-positive Aβ plaques and phospho-tau S262-positive NFTs (**Figure 1**). Many amyloid plaques stained positive for both the Aβ42 and Aβ40 isotype-specific antibodies (**Supplementary Figure S1**); the accumulation of Aβ40-rich parenchymal plaques and cerebral amyloid angiopathy is indicative of more mature, late-stage Aβ deposition in DS [37, 38]. In DS case 4 (36 y), we observed a moderate 4G8-positive Aβ plaque load that was uniformly composed of Aβ42, but not Aβ40, isotypes; moreover, phospho-tau S262-positive tangles were sparse, indicating an early stage of neuropathological burden (**Figure 1** and **Supplementary Figure S1**).

**Figure 1.**
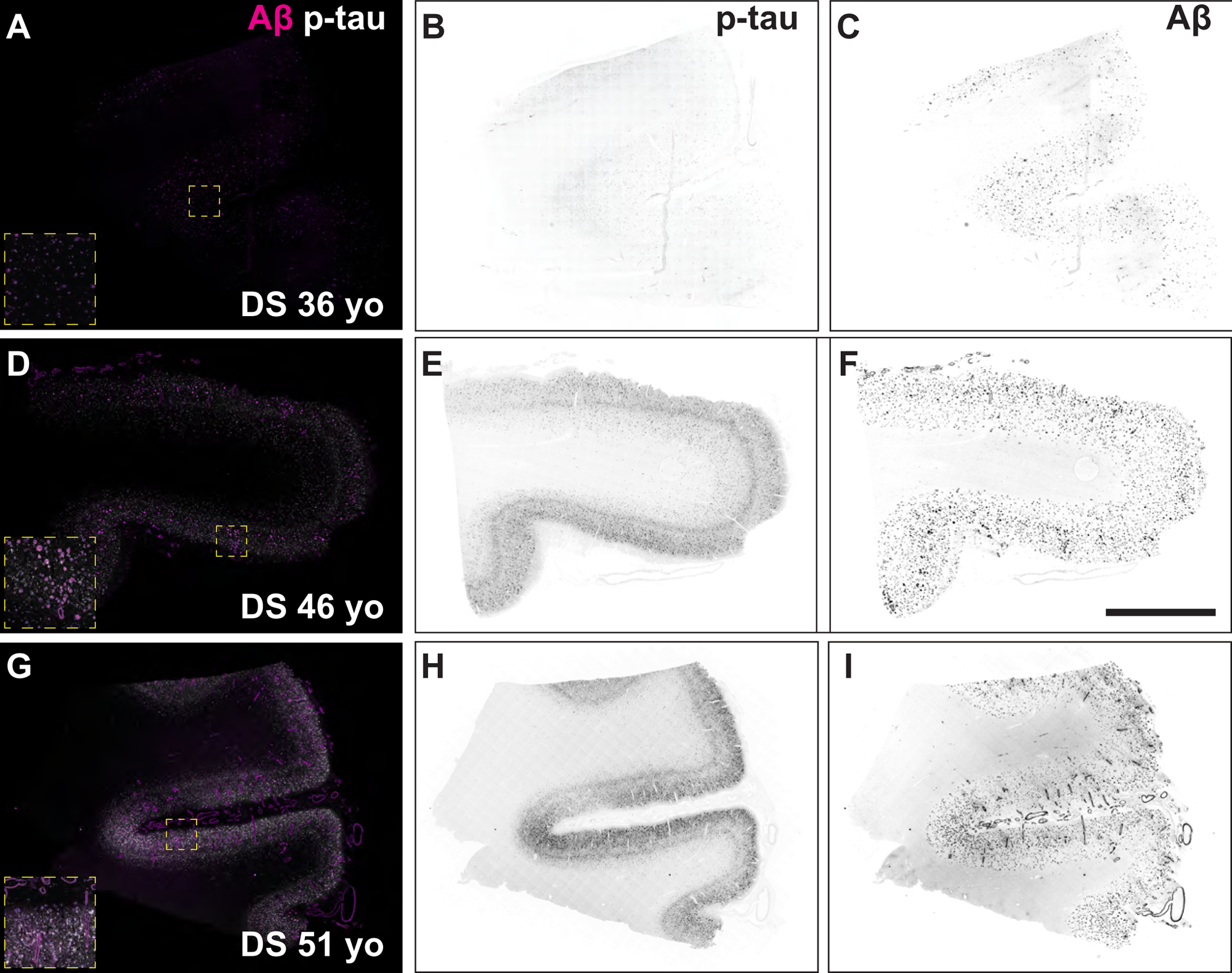
Fluorescence images of immunolabeled Aβ plaques and tau tangles in DS brain samples used for cryo–electron microscopy experiments. Formalin-fixed frontal cortex sections stained with antibodies against total Aβ (magenta) and the phosphorylated tau S262 epitope (white) in DS case 4, 36 years old (A-C), DS case 3, 46 years old (D-F), and DS case 2, 51 years old (G-I). Individual fluorescence channels were converted to grayscale and inverted for phosphorylated tau (B, E, H) and Aβ (C, F, I). Scale bar = 2 mm. Inset dimensions = 0.6 mm^2^.

### In situ assessment of tau tangle conformation in DS

Because older DS brains feature AD neuropathological changes, we postulated that tau conformers should resemble those found in typical late-onset sAD brain samples. Indeed, others have shown that NFTs are positive for three-repeat and four-repeat tau isoforms[39]. However, compared with sAD brains, DS brains exhibit extensive tau pathology in the white matter[40] and produce greater amounts of the three-repeat tau isoform, likely due to other duplicated genes (e.g., *DYRK1A*) on Chr21 that affect the alternative splicing of tau exon 10[39, 41]. Thus, we performed two histological labeling methods in fixed brain slices to assess the conformational nature of tau tangles in DS brains compared with sAD brains. First, we performed immunohistochemistry with a tau AD conformation-specific antibody known as GT-38[42, 43]. Like tau tangles in sAD, we observed that phospho-tau S396-positive NFTs in samples from older and younger DS brains were also positive for GT-38 labeling (**Figure 2A-D**), suggesting that tau filaments in DS may adopt the AD fold.

**Figure 2.**
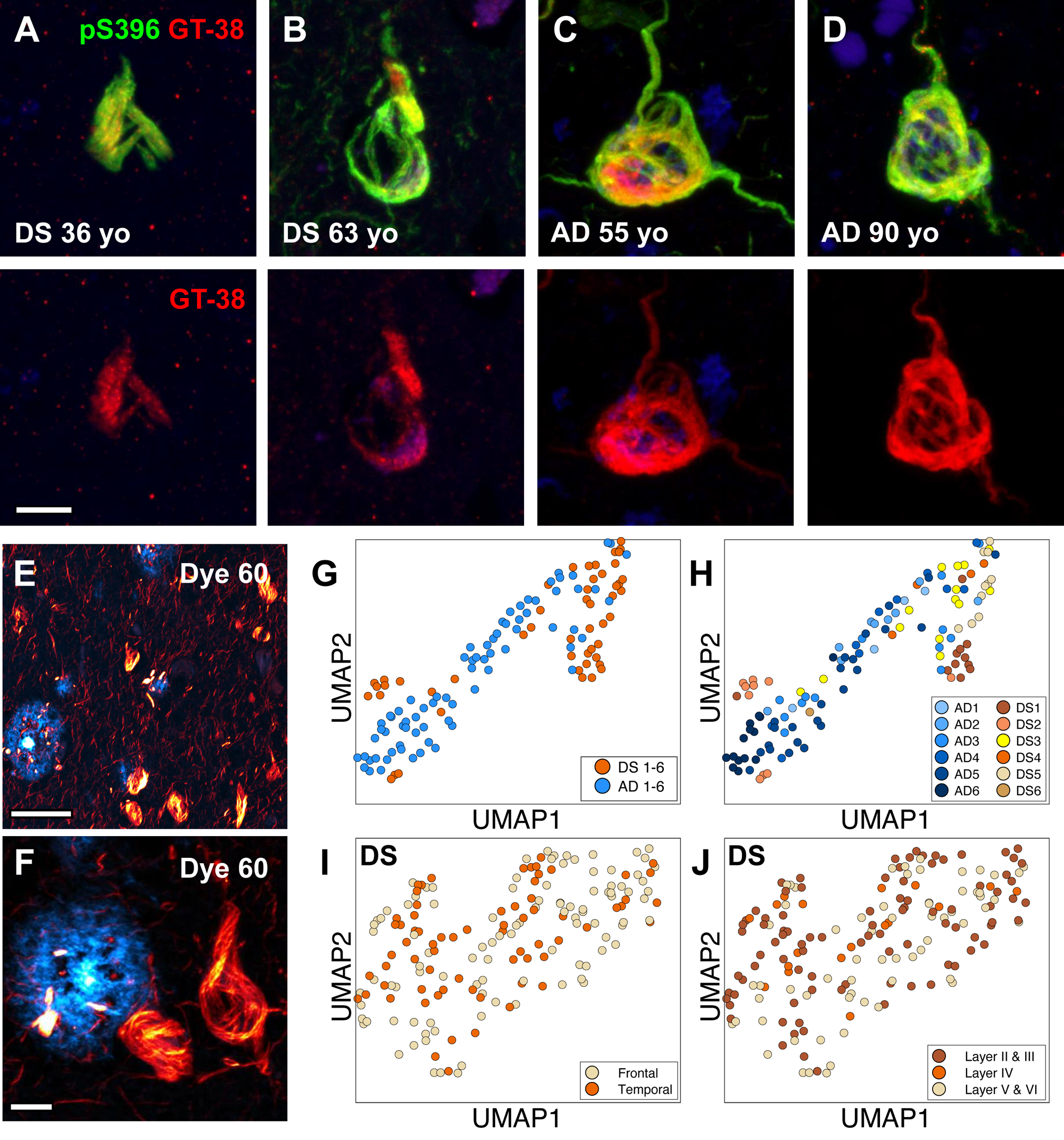
In situ assays show DS tau conformation is similar to tau tangles in sporadic AD. (A-D) Representative confocal images of tau tangles in DS brain fixed slices stained with antibodies targeting the phosphorylated tau S396 epitope (green) and the AD conformation-specific tau fold (GT-38; red). Scale bar = 10 μm. (E, F) Representative 40× low (E) and high (F) zoom confocal images of Aβ plaques (cyan) and tau tangles (red) in DS brain fixed slices stained with dye 60, a novel structure-sensitive dye used for EMBER amyloid-strain discrimination. Scale bar = 50 μm (E); scale bar = 10 μm (F). (G, H) EMBER analysis of tau tangles in brain slices from six DS cases compared with tau tangles in brain slices of six AD cases. EMBER data are plotted in UMAP as all DS cases versus all AD cases (G), or each individual DS case or AD case plotted separately (H). (I, J) EMBER analysis of tau tangles in brain slices from a different set of five DS cases. EMBER data are plotted in UMAP as tau tangles in frontal cortex versus temporal cortex in the same brain (I), or by cortical layers in both frontal and temporal cortices combined (J).

In parallel experiments we used our novel spectral confocal microscopy method, called EMBER imaging, paired with a new structurally sensitive amyloid binding dye, dye 60[20], to assess the conformational heterogeneity of tau tangles in DS and sAD brain slices. Unlike traditional fluorescent amyloid dyes, like thioflavin S, Congo red, or their derivatives, our dye 60 exhibits robust differences in the excitation and emission spectra when bound to Aβ plaques versus tau tangles, thus making for facile spectral separation and morphological distinction between these two protein deposits (**Figure 2E, F**). We previously demonstrated EMBER could robustly discriminate tau deposits in sAD and Pick’s disease[20]. Here, we stained six DS samples (including cases 1, 2, and 4) and six sAD samples. Following our published protocol (see Methods), we collected and analyzed EMBER data for tau tangles only and presented the results in a UMAP plot. The EMBER signature of tau tangles in a DS brain clustered tightly with tau tangles in a sAD brain (**Figure 2G, H**), suggesting that the DS tau tangle conformation may be like those in AD. Next, we examined the degree of conformational homogeneity between and within brain regions in the same DS cases. We obtained a new set of fixed postmortem tissues with slices from both the frontal cortex and temporal cortex from five individuals with DS and advanced AD neuropathology. We performed EMBER imaging and observed that the conformational signature of tau tangles was similar in the frontal and temporal cortices and across all cortical layers in those regions (**Figure 2I, J**). To look more broadly at tau conformers in DS and AD, we performed sarkosyl extraction of insoluble tau from fresh-frozen frontal cortex samples (**Supplementary Table S2**) and used a HTRF assay to measure aggregated tau species (**Supplementary Figure S2A, B**) to select a subset of well-matched samples for Western blot (**Supplementary Figure S2C–F**). We performed limited proteolysis with trypsin followed by Western blot analysis on undigested and digested sarkosyl-insoluble tau extracts using a panel of antibodies targeting different tau epitopes. We observed no discernible differences in the antibody staining or banding pattern of insoluble tau from individuals with DS compared to AD patients (**Supplementary Figure S2**). Taken together, these data corroborate the notion that tau filaments from DS are likely similar to the AD tau fold.

### Cryo-EM structures reveal tau filaments adopt AD conformations in DS

We sought to determine structures of isolated tau filaments from DS cases 1 to 4 (**Supplementary Table S1**) by cryo-EM. Sarkosyl-insoluble material was extracted from frontal cortex tissue essentially as described[13, 21] (see Methods) and initially assessed by negative-stain EM (**Supplementary Figure S3**). PHF-like filaments were readily identified in cases 1 to 3; however, very few fibrils were observed for case 4. We note this was expected based on the younger age of the individual (36 y) and the presence of fewer phospho-tau S262-positive tangles compared with other cases (**Figure 1**). More narrow filaments were also identified in some cryo-EM micrograph images; these were attributed to Aβ, as has been recently described for DS[44], and were not further characterized in this study given their low prevalence relative to tau (**Supplementary Figure S3B**). For case 1, cryo-EM 2D and 3D analysis revealed the presence of both PHFs and SFs, totaling approximately 76% and 21% of the selected data, respectively (**Figure 3A**). Cryo-EM helical reconstructions using RELION[26, 27] resulted in well-resolved PHF and SF final maps at 2.7-Å and 2.9-Å resolution, respectively (**Figure 3B**, **Supplementary Figure S4**, and **Supplementary Table S4)**. The docked atomic models for tau residues G303 to E380 were in good agreement with the map and confirmed the tau conformations are identical to the AD PHF and SF structures (**Figure 3C**, **Supplementary Figure S4**, and **Supplementary Table S4**). Indeed, alignment to the AD structures showed no substantial differences, with an overall α-carbon root mean square deviation equal to 0.5 Å and 0.6 Å for the PHF and SF, respectively (**Supplementary Figure S4**). As with AD, the filaments adopted a characteristic *C*-shaped fold comprising a β-helix turn (residues 337-356) and an antiparallel protofilament interaction interface across the _332_PGGGQ_336_ motif for the PHF and residues 317-324 and 312-321 for the SF.

**Figure 3.**
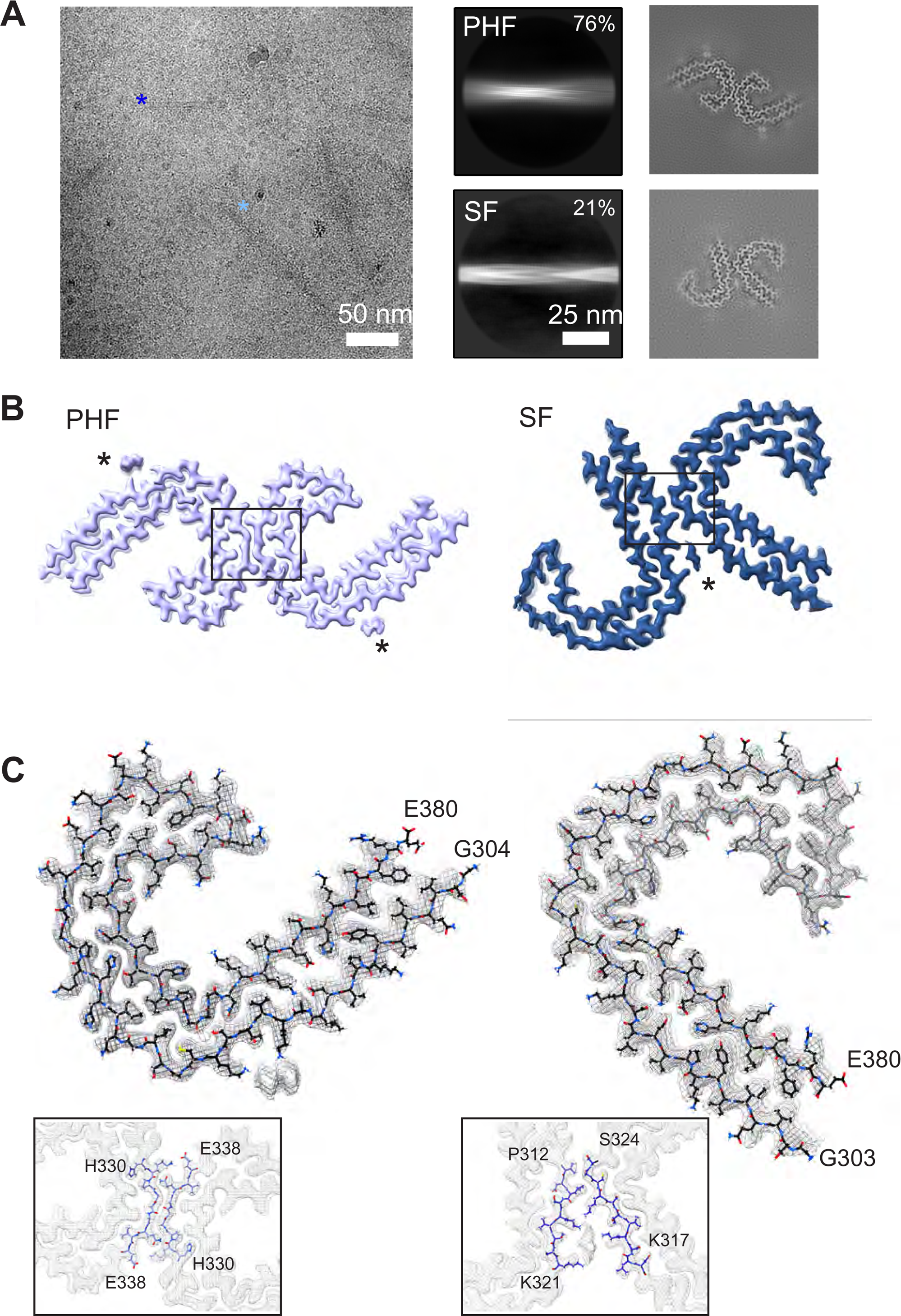
Cryo-EM structures of tau PHF and SF conformations from DS case 1. (A) Representative electron micrograph of amyloid filaments from the frontal cortex of DS case 1 showing PHF and SF segments (left image; light and dark blue asterisks, respectively) and corresponding 2D class averages (middle images) and cross-section projection views of the conformations (right images). (B) Final cryo-EM density maps of the PHF and SF at 2.7-Å and 2.9-Å resolution, respectively. Asterisks indicate extra densities previously observed in structures from AD, and boxes show each protofilament interface, enlarged in (C). (C) Single protofilament cryo-EM map (mesh) with the fitted atomic model for the PHF (left) and SF (right), including tau residues G304 for PHF (G303 for SF) to E380 and an enlarged view of the protofilament interface.

Considering the very low abundance of tau filaments and scarcity of tissue for case 4, we devised a scheme to enrich for tau filaments on cryo-EM grids prior to vitrification using an established affinity-grid method (**Figure 4A**)[23]. For this procedure, single-layer GO was applied to QUANTIFOIL grids, functionalized with a PEG maleimide, and coupled to protein G for binding to the AT8 phospho-tau (S202 and T205) antibody (see Methods). This GO-AT8 affinity-grid method was tested initially for capturing tau filaments purified from case 2 samples, which exhibited robust NFTs. In control experiments, in which protein G was not added to the affinity-grid assembly procedure, few to no filaments were observed in cryo-EM images (**Figure 4B**). We note this procedure involved wash steps after sample application to reduce background contaminants such as ferritin[13], which can impact filament selection and refinement. Thus, the absence of filaments indicated successful removal of unbound material relative to typical vitrification procedures. Conversely, when AT8 was included in the grid preparation, we observed extensive tau filaments in cryo-EM images, although some background contaminants remained (**Figure 4C**). Together, these data indicate our GO-AT8 affinity-grid method successfully enriched for tau filaments in an AT8-dependent manner.

**Figure 4.**
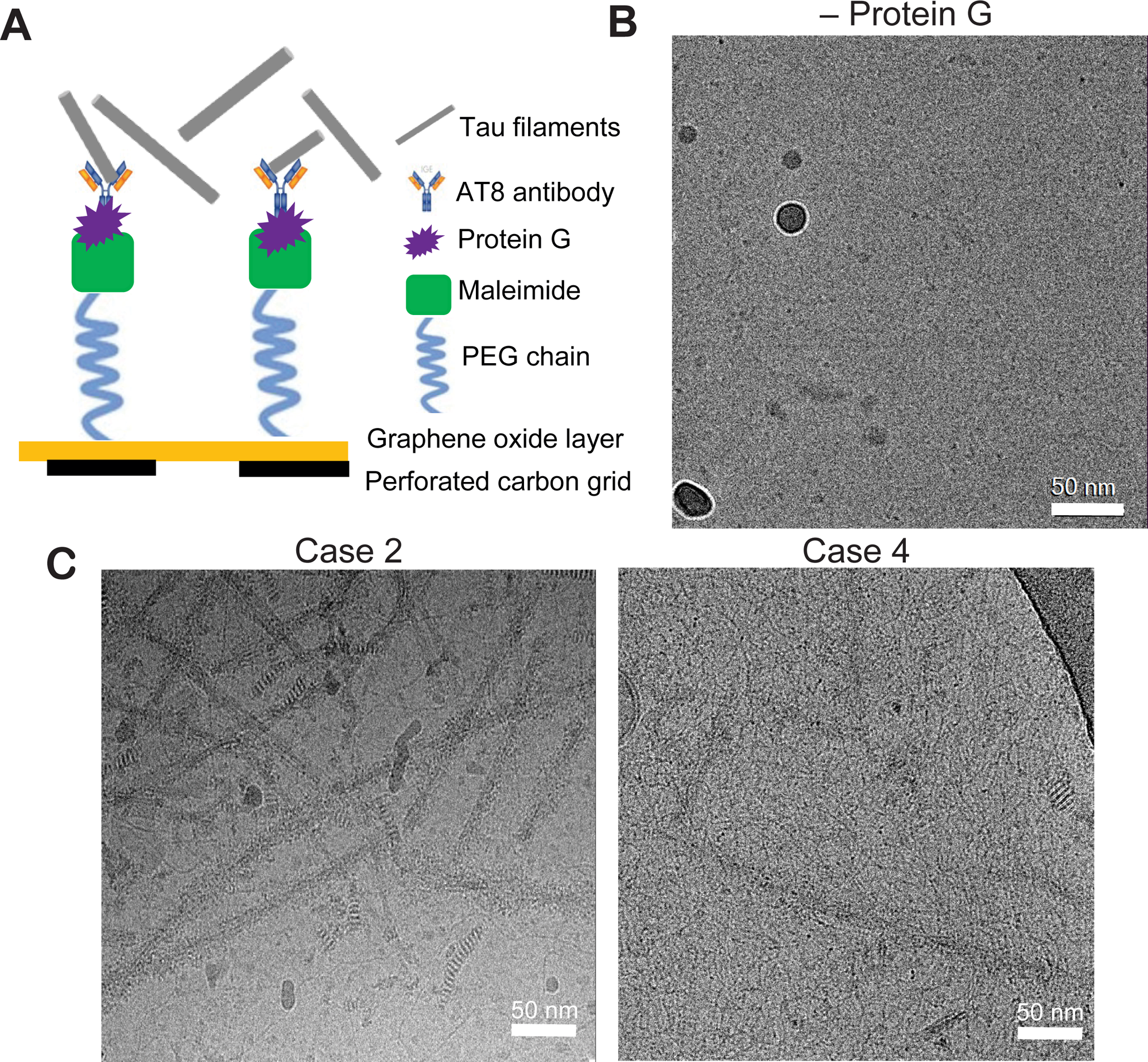
GO-AT8 antibody affinity-grid assembly and isolation of tau filaments for cryo-EM. (A) Schematic showing the affinity-grid assembly procedure for the attachment of AT8 anti-tau antibodies to the GO grid surface. (B) Control cryo-EM micrograph image for which protein G was not included in the affinity-grid assembly prior to incubation with tau filaments and vitrification. (C) Representative cryo-EM micrograph images following GO-AT8 affinity-grid isolation of tau filaments from DS case 2 (left) and DS case 4 (right). Fewer filaments are observed in case 4 due to the low overall abundance of tau deposits.

Next, cryo-EM datasets were collected for tau filaments isolated from DS cases 2 to 4, with GO-AT8 affinity grids being used for cases 2 and 4. The resulting cryo-EM reconstructions showed all cases contained predominantly AD PHF conformations, while cases 2 and 3 also exhibited substantial SFs, similar to case 1 (**Figure 5**). Notably, structure determination for case 4 remained limited by the low abundance of isolated filaments, resulting in a final resolution of 5 Å for the case 4 PHF. However, molecular models for cases 2 and 4 confirmed PHF and SF conformations were resolved using the GO-AT8 grids (**Supplementary Figure S5)**. Interestingly, a small dataset (approximately 17,000 segments) was classified and refined to a low-resolution (>7 Å) map that appears similar to the CTE type II fold found in CTE [16] and SSPE [45]. However, the map resolution and quality were not sufficient to build an atomic model and confirm the precise tau filament conformation. Although some overfitting is apparent in the 2D slice view (**Figure 5**), we note that the reference-free 2D class averages selected for the reconstruction appear distinct from the PHF averages and are similar to averages reported for CTE (Supplementary Figure S5) [16]. Overall, for the four cases we studied, these data identify that the tau AD PHF fold predominates across a wide age range of individuals with DS. The SF fold was also present at lower levels but is comparable to what has been previously observed by cryo-EM[13]. Therefore, both the structural conformation of tau filaments and the relative presence of PHFs and SFs appear identical to those in AD for older individuals with DS with a higher burden of tau deposits, supporting the general AD pathology that has been reported for DS[3].

**Figure 5.**
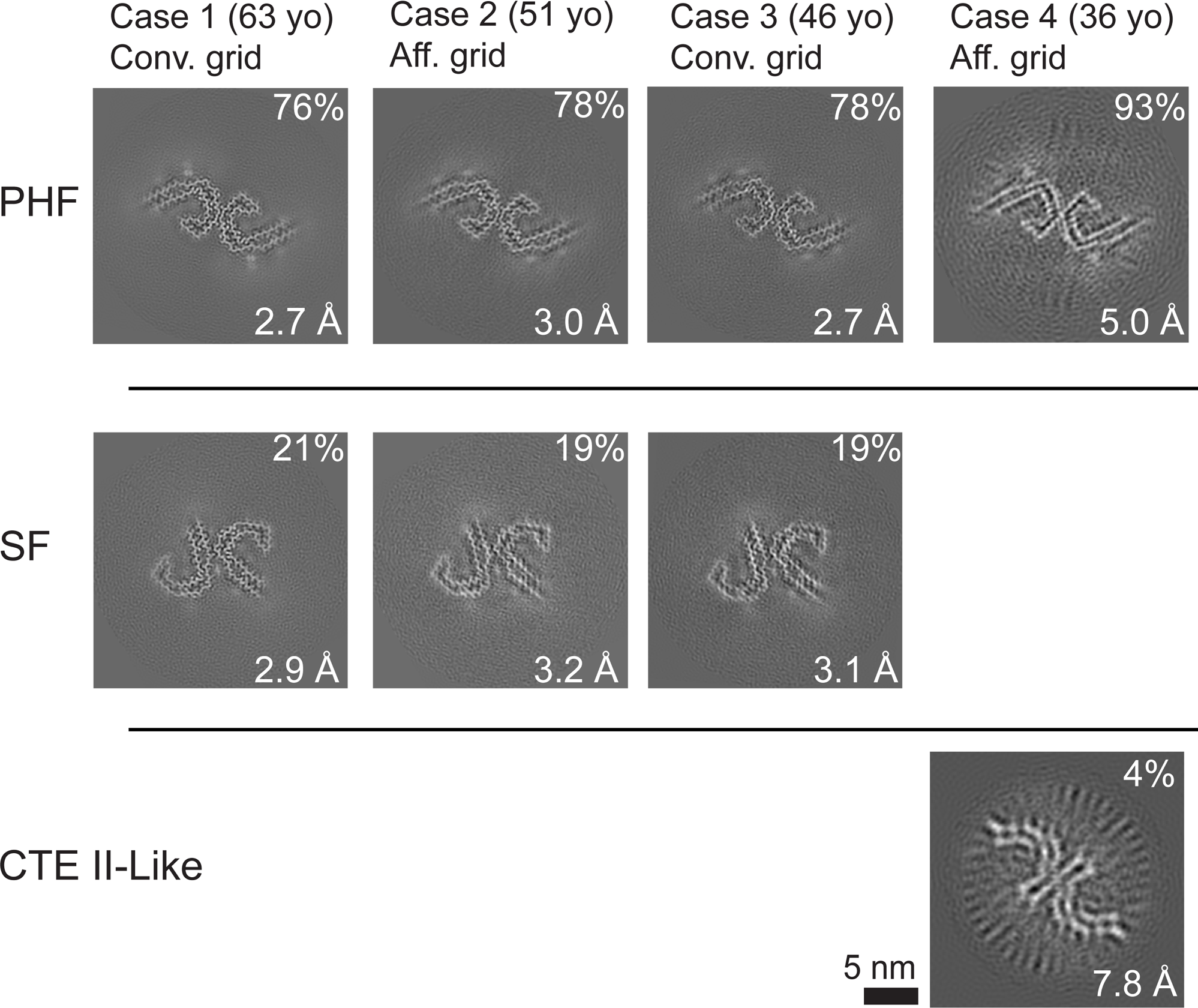
PHF and SF cross-section views of the tau filament cryo–electron microscopy structures determined from four DS cases. The relative percentage of the segments for the reconstructions is indicated with the final resolution and grid preparation method. In the 36 years old case, a minor percentage refined to show a conformation similar to CTE type II filaments (indicated). However, an atomic model was unable to be determined due to the low resolution and small dataset size.

## DISCUSSION

Cryo-EM structural studies have revealed that tau adopts a myriad of polymorphic folds that are associated with distinct etiological forms of primary and secondary tauopathy diseases[18]. In individuals with DS, clinical biomarkers of tau pathology progress along a similar trajectory as for people with AD—just decades sooner[8]. This early-onset AD has long been attributed to the triplication of *APP* and exacerbated deposition of Aβ neuritic plaques[3, 4]. Moreover, isolated insoluble tau[46, 47] and NFTs [3, 48] that arise in individuals with DS share similar biochemical and morphological properties to the disease form of tau seen in AD. However, structures of tau filaments from individuals with DS have not previously been determined. Thus, the specific tau fold that develops in DS has been unknown; a high-resolution structure of tau in DS would bear great significance for therapeutic development. Our novel workflow of *in situ* conformational assays and cryo-EM methods revealed that DS tau adopts the AD folds of PHFs and SFs.

We previously developed EMBER imaging with structure-sensitive dyes to visualize conformational heterogeneity of intracellular tau deposits in situ and identified robust conformational discrimination between individual AD NFTs and Pick bodies at the cellular level[20]. We therefore used EMBER imaging to screen and predict the conformational status of tau deposits in candidate brain samples and, indeed, demonstrated a robust overlap of conformational signatures for NFTs in AD and DS samples (**Figure 2G, H**). In AD, tau NFTs accumulate predominantly in the deeper cortical layers V/VI[49]; in contrast, tau NFT accumulation is more widespread in DS, involving the cortical layers II through VI[48, 50]. In a small cohort of well-matched DS cases, we found that the EMBER-based conformational signature of DS tau NFTs was homogeneous across all cortical layers in the frontal and temporal cortices (**Figure 2I, J**). These data suggest that (a) the same AD tau folds should be found in the temporal cortex of individuals with DS and that (b) cortical neurons in layers II/III in DS give rise to NFTs composed of tau filaments with the same AD tau fold as neurons in layers V/VI in DS or AD. Considering the overexpression in DS of more than 250 protein-coding genes on Chr21, it is remarkable that the DS tau fold is identical to those in AD cases and that the tau NFT conformations in DS appear homogeneous within and between DS cases.

It has been postulated that the distinct CTE tau fold[16], also found in SSPE[45] and amyotrophic lateral sclerosis and parkinsonism-dementia complex (ALS-PDC)[51], derives from tau NFT deposition predominately in the upper cortical layers II/III as a consequence of distinct disease mechanisms involving chronic neuroinflammation and/or environmental factors[52–55]. In contrast, we show that DS tau NFTs in layers II/III exhibit a similar EMBER signature as NFTs in layers V/VI, suggesting that any putative microenvironmental differences specific to each cortical layer are not sufficient to drive the formation of CTE tau folds versus AD tau folds. Hence, it is reasonable to posit that a distinct neuroinflammatory state shared between CTE, SSPE, and ALS-PDC drives the formation of a common CTE tau fold that is unique from AD tau PHFs and SFs. Nevertheless, trisomy of Chr21 leads to a heightened state of neuroinflammation that is not typical of AD. In DS, there is triplication of four interferon receptor genes on Chr21, causing a lifelong, systemic type 1 interferonopathy phenotype (a chronic state of low-grade inflammation in all organs)[56, 57]. In the brain, proinflammatory cytokine profiles and aberrant microglial activation states are more pronounced in DS compared with AD[58]; in later stages of DS, a robust neuroinflammatory phenotype emerges, including a prominent dystrophic and rod-shaped microglial phenotype that aligns with neurons forming tau tangles[59, 60]. Therefore, it is somewhat surprising that the AD tau conformation predominates in the DS brain despite the overt differences in the biochemical microenvironment. Thus, we conclude that the Aβ pathology, and its associated cellular consequences, must be the major driver of the PHF and SF tau conformations no matter the etiological form of AD.

In DS, overt deposition of Aβ plaques and tau NFTs appears in the second and third decade of life, respectively; however, cognitive impairment and dementia do not emerge until the fifth or sixth decade of life[2]. The field now has a clear view of the spectrum of distinct tau folds present in late stages of disease, but the extent of conformational heterogeneity at disease onset remains unknown. To increase the potential for translational success, it is important to determine the structures of tau filaments from the very early asymptomatic stages of disease. Critical to the characterization of tau filaments from the younger (36 y) individual with no documented dementia was our implementation of GO-AT8 affinity grids to isolate tau filaments from a very sparse sample (**Figure 1**). This procedure enabled small-scale affinity purification from an endogenous source, an advantageous step for future potential structural efforts targeting specific brain regions, early disease stages, or other pathologies with reduced tau deposits. We additionally found this procedure to be beneficial in reducing background contaminants, such as ferritin, which can impact classification and refinement. Unfortunately, despite these advances, we were unable to complete high-resolution structures from the 36-year-old case; thus, dataset size and filament abundance remains a challenge. Nonetheless, our structure determined to 5.0-Å resolution was sufficient to establish the AD PHF fold. Therefore, further optimizations were not pursued.

Strikingly, with the GO-AT8 affinity-grid method, we were able to identify that 4% of tau filaments from case 4 appeared to have the type II CTE fold, also found in SSPE and ALS-PDC[16, 45, 51]. However, due to the extreme rarity of these tau filaments, we only achieved a 7.8-Å map and thus cannot conclusively determine the fold to high resolution. Nevertheless, if we assume that more data would confirm there is a minor population of type II CTE tau filaments in case 4, how do we reconcile that the other three cases with advanced AD pathology and abundant tau filaments do not have this morphology? One explanation could be that case 4 also had a well-documented history of epilepsy and the cause of death was respiratory failure. Notably, epilepsy (in non-DS young adults) is associated with the accumulation of tau histopathology[61–63], and neuroinflammatory pathways are activated by and exacerbate epileptic seizures[64]. Thus, we speculate that the neurodegenerative and neuroinflammatory processes in the epileptic brain may precipitate conditions that yield the type II CTE fold. As individuals with DS live longer into late middle age, the prevalence of comorbid epilepsy is estimated to reach upward of 50% of the DS population with AD[65]. Future cryo-EM studies on tau filaments from greater numbers of individuals with DS and comorbid epilepsy may identify cases with a heterogeneous mix of AD tau and CTE tau folds, consistent with a prior study showing multiple tau polymorphs identified in a single primary tauopathy case[18].

Overall, our findings are in good agreement with a recent study from Fernandez and colleagues[44], who published their work at the time our manuscript was submitted for peer review. Juxtaposed with our cryo-EM and EMBER imaging data in a larger DS cohort, there is strong evidence that tau filaments adopt the AD PHF fold early in the neuropathogenesis of DS and persists as the predominant tau fold in older individuals with DS. While Aβ filament structures were not the focus of our study here, Fernandez et al. determined the cryo-EM structures of Aβ42 filaments that are identical to those found in sporadic and inherited AD[21] as well as discovering two new Aβ40 filaments (types IIIa and IIIb)[44]. While these novel Aβ40 filaments were not yet detected in sporadic or inherited AD samples, it will be interesting to see if this observation is related to the increased Aβ40-rich cerebral amyloid angiopathy that can be more prominent in DS compared to AD[66]. Alternatively, the novel Aβ40 filaments could be linked to the presence of unique parenchymal plaque type in DS not found in AD, consistent with our previous data showing that a subset of older individuals with DS featured a distinct conformational signature of Aβ plaques compared to sporadic AD patients[36]. Cryo-EM structural studies of Aβ filaments should be expanded to a larger DS cohort including samples from young individuals with DS. Deciphering the extent of Aβ filament heterogeneity across the lifespan of individuals with DS may have important implications for drug discovery.

Because modern clinical imaging and fluid biomarkers have revealed the trajectory of AD progression in individuals with DS[8–10], there has been growing consideration to include individuals with DS in clinical trials for therapeutics developed for AD. The urgency to do so has been tempered by the fact that the mechanisms of action and/or safety profiles of candidate drugs may be different in individuals with DS, despite the shared presence of Aβ plaques and tau tangles. Indeed, the first clinical trial (NCT05462106) testing an Aβ vaccine in participants with DS is underway[67], and there may be future trials testing Aβ immunotherapies that are already showing promising results in individuals with sAD. However, the formation of tau tangles after the deposition of Aβ plaques is much more rapid in DS than in sAD[68, 69]; thus, effective therapeutics targeting pathological tau in DS will be required due to the condensed trajectory of AD pathogenesis. Given the intracellular localization of tau NFTs, immunotherapy approaches are unlikely to succeed, emphasizing the need to develop small-molecule therapeutics. By solving the structure of tau filaments in DS, we now have a molecular framework to apply existing or design new conformation-specific small-molecule drugs or diagnostics that target the AD tau fold. For example, we expect that the cleft of the PHF fold in DS bears an identical pocket for stacked small-molecule binding, as we have demonstrated with the positron emission tomography ligand GTP1[70] and as Kunach et al. demonstrated with the positron emission tomography ligand MK-6240[71] in tau PHFs from sAD. Based on this novel binding mode, new *in silico* docking methods[72] may predict small molecules with superior properties for development as next-generation diagnostic ligands or therapeutic inhibitors targeting the tau fold in DS and AD.

## ACKNOWLEDGMENTS

This work was supported by grants from the National Institutes of Health (NIH; P01AG002132 to S.B.P., C.C., and D.R.S.; RF1NS133651 to C.C. and D.R.S.), Henry M. Jackson Foundation (HU0001-21-2-065 subaward 5802 to S.B.P.), and Rainwater Charitable Foundation (A138221 to D.R.S.). H.Y. received postdoctoral fellowship funding from the BrightFocus Foundation (A2020039F). Tissue samples were supplied by the NIH NeuroBioBank (via the University of Maryland); the August Pi i Sunyer Biomedical Research Institute Biobank (Barcelona, Spain); the London Neurodegenerative Diseases Brain Bank (King’s College London, UK), which receives funding from the Medical Research Council, UK, and the Brains for Dementia Research project (jointly funded by the Alzheimer’s Society and Alzheimer’s Research UK); Autistica (London, UK); the National Institute for Health and Care Research Oxford Biomedical Research Centre; and Professor Elizabeth Head and the University of California, Irvine, Alzheimer’s Disease Research Center (Irvine, CA), which is funded by the NIH National Institute on Aging (grant P30AG066519).

## AUTHOR CONTRIBUTIONS

U.G., E.T., D.R.S., and C.C. designed the research; U.G., E.T., H.Y., M.S., C.D.C., F.W., and C.C. performed the experiments; U.G., E.T., H.Y., G.E.M., S.B.P., D.R.S., and C.C. analyzed the data; and U.G., E.T., D.R.S., and C.C. wrote the paper. All authors read and approved the final manuscript.

## AVAILABILITY OF DATA & MATERIALS

Cryo-EM maps will be available in the Electron Microscopy Data Bank (EMDB) upon publication. Corresponding refined atomic models will be deposited in the Protein Data Bank (PDB) upon publication. Please address requests for materials to the corresponding authors.

## SUPPLEMENTARY INFORMATION

**Supplementary Figure S1.**
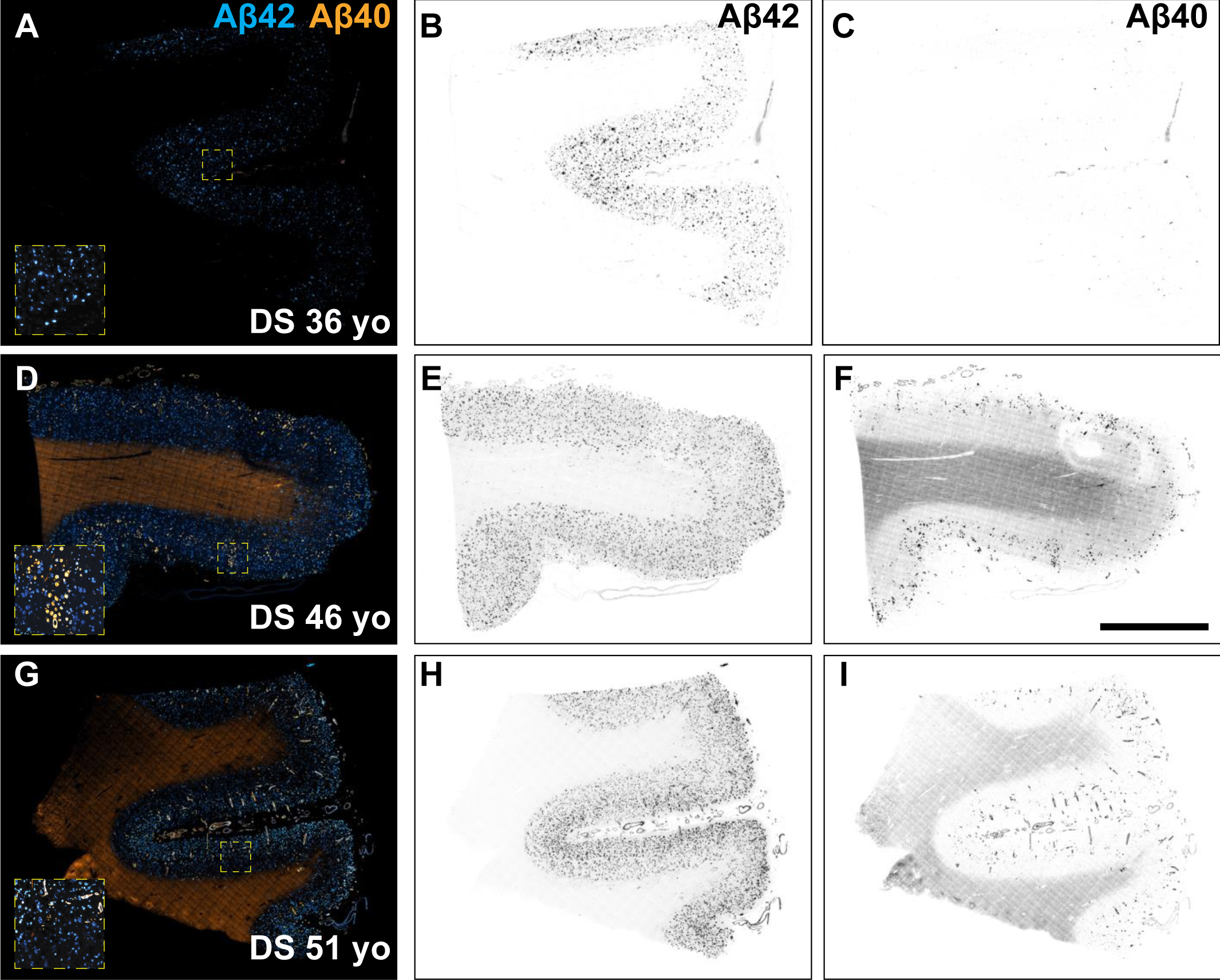
Fluorescence images of immunolabeled Aβ42 and Aβ40 plaques in DS brain samples used for cryo–electron microscopy experiments. Formalin-fixed frontal cortex sections stained with Aβ isotype-specific antibodies against C-terminal-length isoforms Aβ42 (cyan) and Aβ40 (orange) in DS case 4, 36 years old (A-C), DS case 3, 46 years old (D-F), and DS case 2, 51 years old (G-I). Individual fluorescence channels were converted to grayscale and inverted for Aβ42 (B, E, H) and Aβ40 (C, F, I). Scale bar = 2 mm. Inset dimensions = 0.6 mm^2^.

**Supplementary Figure S2.**
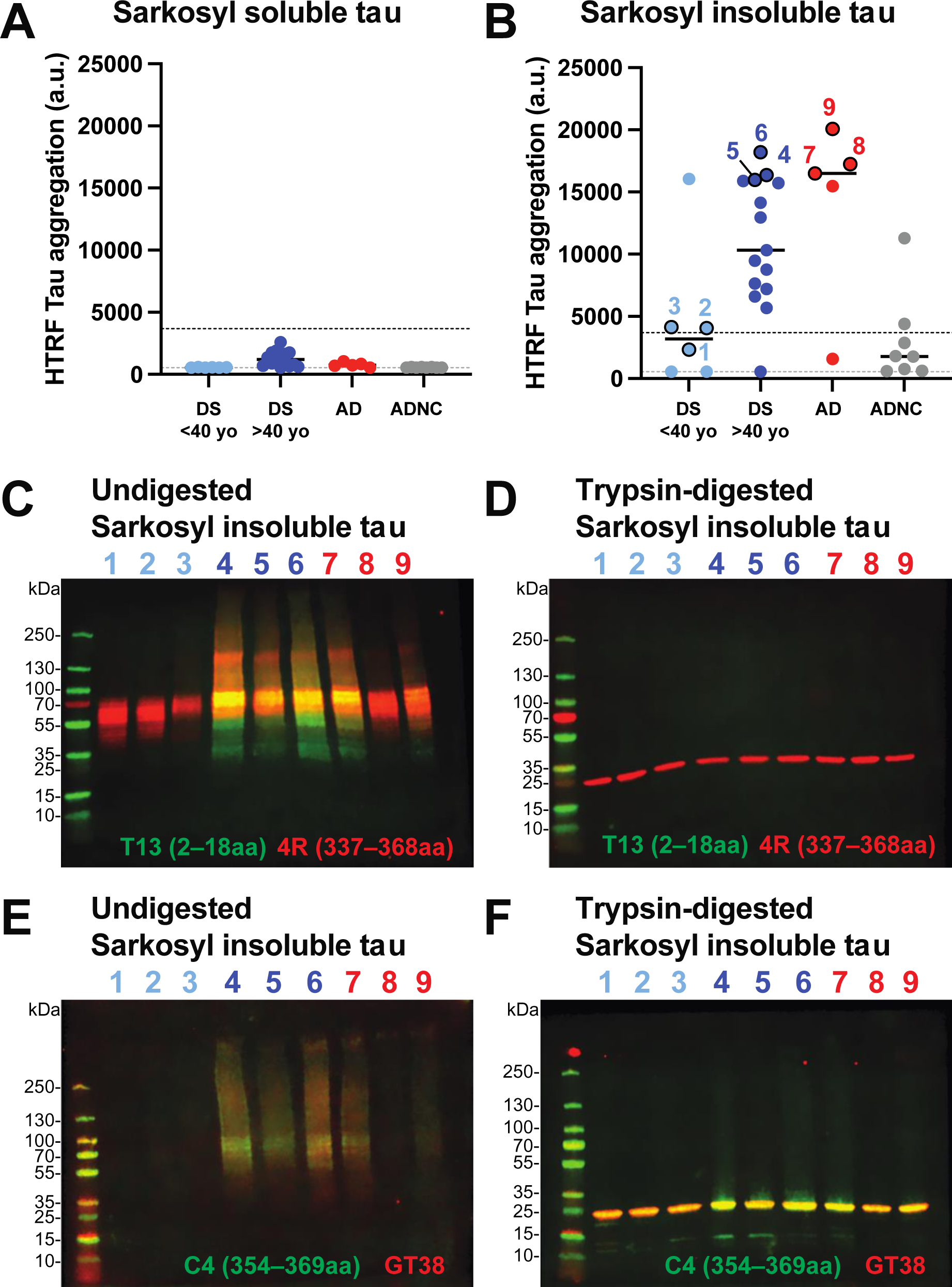
Biochemical characterization of sarkosyl-insoluble tau from DS and AD frontal cortex. (A and B) Quantification of aggregated tau in sarkosyl-soluble and sarkosyl-insoluble extracts from frozen frontal cortex of DS (<40 years old and >40 years old), AD, and ADNC (Alzheimer’s disease neuropathologic change; individuals with no cognitive impairment but have presence of modest AD neuropathology). HTRF assay positive control and negative control values are represented by the upper and lower dotted y-axis lines, respectively. HTRF Tau aggregation assay shows only marginal aggregated tau in the sarkosyl-soluble fraction for DS (<40 yo), AD, and ADNC with the exception of DS (>40 yo) group showing modest levels of aggregated tau for some samples (A). HTRF Tau aggregation assay shows robust levels of aggregated tau in the sarkosyl-insoluble fraction for most of the DS (>40 yo) and AD samples; modest levels of aggregated tau were detected for a subset of DS (<40 yo) and ADNC samples (B). Based on the HTRF Tau aggregation values in the sarkosyl-insoluble fraction, we selected a subset of samples for Western blot as indicated by the numbers on panel B. (C–F). Western blot analysis of undigested and trypsin-digested sarkosyl-insoluble tau. (C and D) T13 and R4 antibodies and (E and F) C4 and GT38 antibodies. The tau protein segments recognized by the T13, 4R, and C4 antibodies are specified in parentheses. The undigested tau bands appear primarily at ∼60 kD, and the trypsin-digested tau bands at 25 kD. Trypsin cleaves the *N*-terminal unstructured domain of tau, resulting in the absence of signal for the T13 antibody (residues 2–18) as shown in panel D. Trypsin is unable to cleave the fibril core (residues 303–380), thus signals for the 4R antibody (residues 337-368) and the C4 antibody (residues 354-369) are present in the trypsin-resistant band at 25 kD (D and F). The GT38 antibody, which is specific to the AD tau conformation (exact binding epitope unknown), also binds to the trypsin-resistant fibril core observed in al DS and AD cases indicating similar epitope and conformation is present across all samples.

**Supplementary Figure S3.**
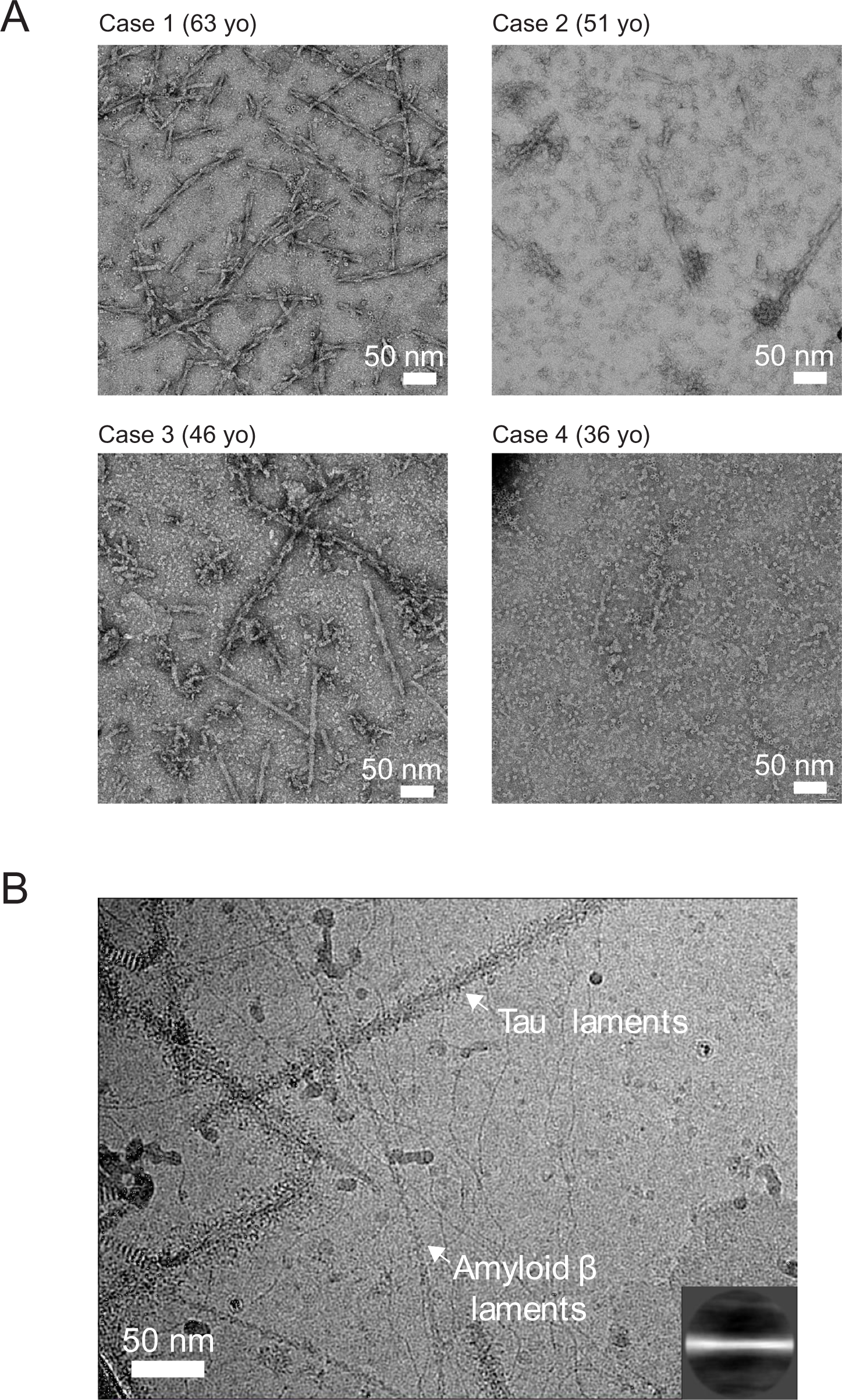
Electron microscopy images of filaments from sarkosyl-insoluble fractions from the frontal cortex of Down syndrome. (A) Negative stain images from the four cases (indicated) showing fibrils and other background contaminants. (B) Cryo-EM micrograph image showing tau filaments and filaments attributed to amyloid beta. A 2D class average of a small dataset that shows features similar to amyloid beta filaments is shown (inset).

**Supplementary Figure S4.**
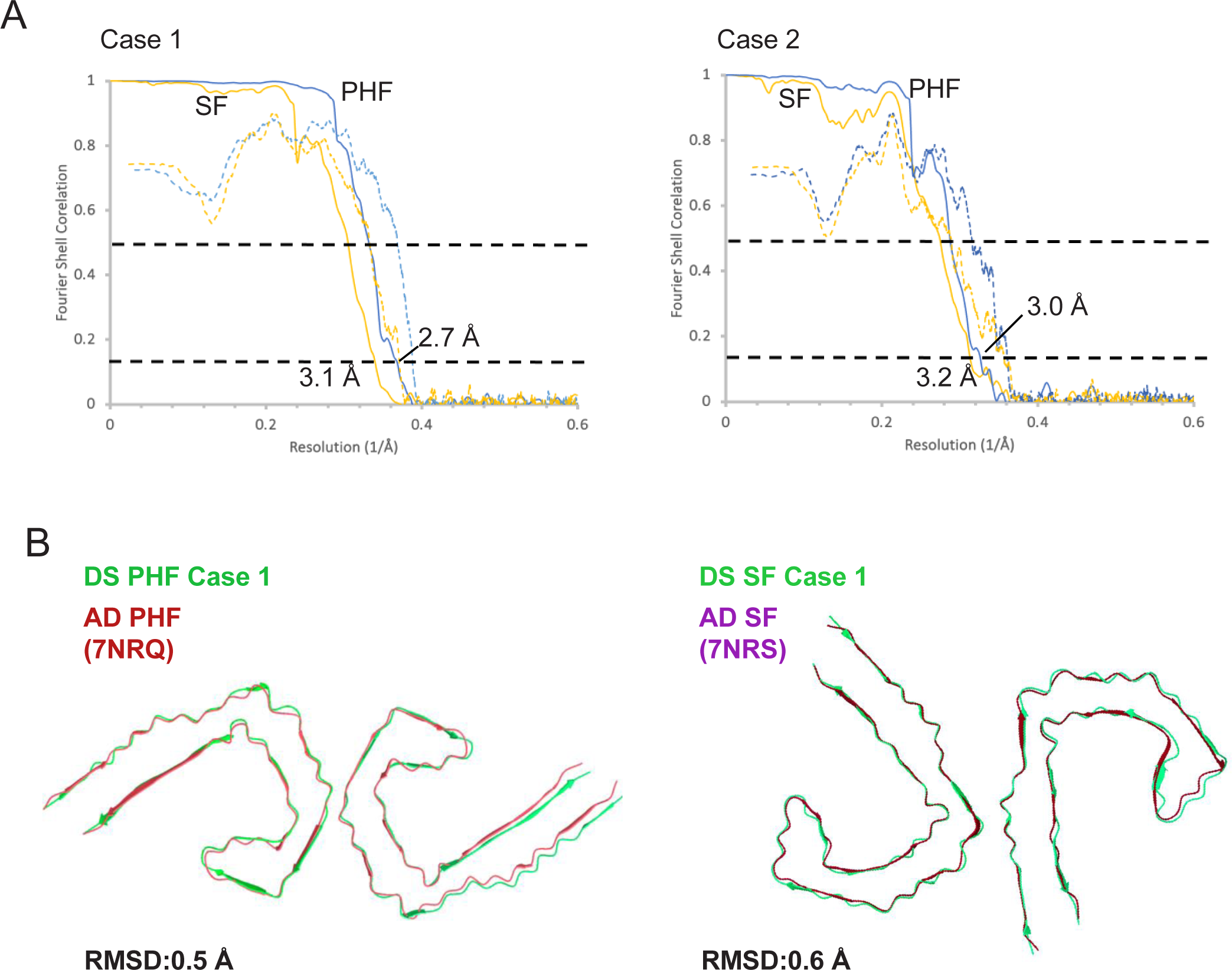
Cryo–electron microscopy structure analysis. (A) FSC curves of two independently refined half-maps for the PHF (blue) and SF (orange) showing resolution at FSC = 0.143 and corresponding map versus model curves (dashed) and FSC = 0.5 line. (B) Overlay and α-carbon RMSD values for the PHF and SF structures from case 1 (green) compared with published structures (Protein Data Bank: 7NRQ [red] and 7NRS [purple], respectively).

**Supplementary Figure S5.**
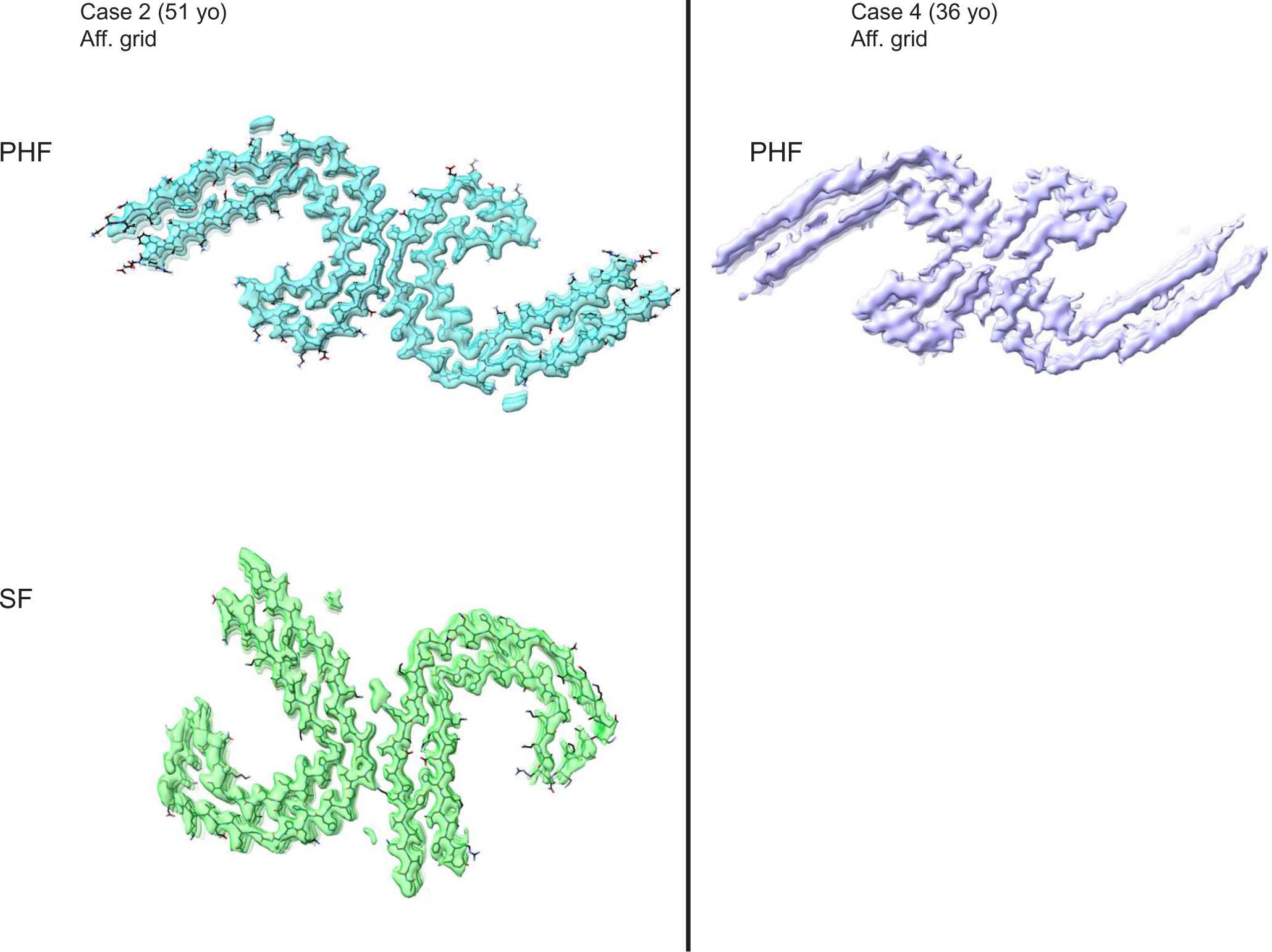
Final cryo–electron microscopy refined maps of tau PHF and SF from data acquired using GO-AT8 affinity grids to isolate tau filaments. For DS case 2, the PHF structure was solved to 3.0 Å and the SF structure was solved to 3.2 Å resolution (left); for DS case 4, the PHF structure was solved to 5 Å resolution (right).

**Supplementary Figure S6.**
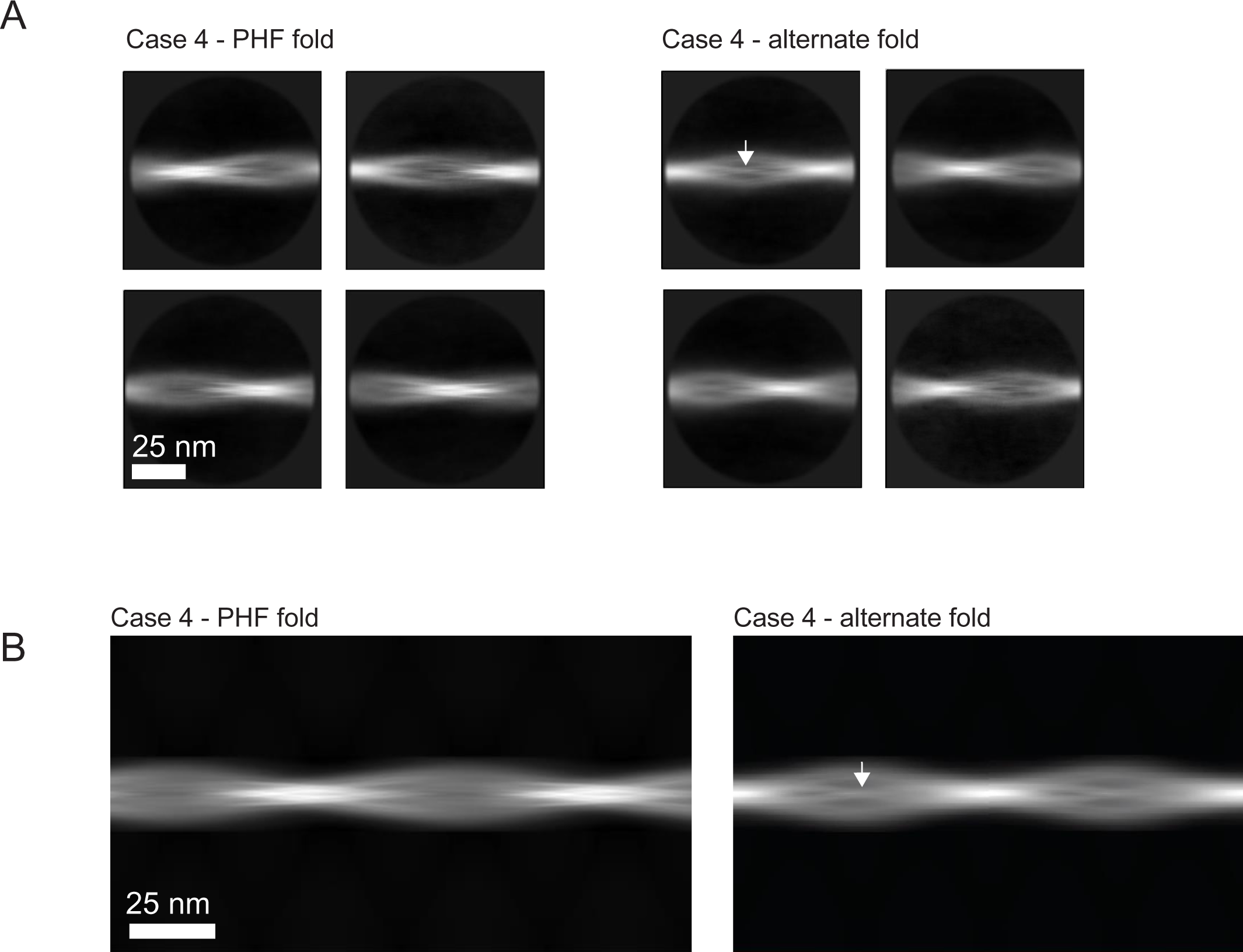
Comparison of 2D averages observed for tau filaments from DS case 4. (A) 2D averages comparing a PHF class and one identified to be in an alternate conformation (CTE) [1] based on distinguishing density in the middle the wide portion of the crossover (arrow). (B) 2D averages generated using relion_helix_toolbox by stitching to show full crossover for comparison between the PHF and alternate fold. The central line of density unique to the alternate fold is indicated (arrow). 1. Falcon B, Zivanov J, Zhang W, et al (2019) Novel tau filament fold in chronic traumatic encephalopathy encloses hydrophobic molecules. Nature 568:420–423. https://doi.org/10.1038/s41586-019-1026-5.

**Supplementary Table S1.** Fresh-frozen postmortem donor tissues used for filament extraction and cryo–electron microscopy. Abbreviations: Aβ, amyloid-β; APOE, apolipoprotein E; BCN, Barcelona; CERAD, Consortium to Establish a Registry for Alzheimer’s Disease; DxA, density × area; F, female; M, male; ND, not determined; NFT, neurofibrillary tangle; UMD, University of Maryland; Xtau, custom measurement of immunostained neurofibrillary tau tangles in fixed medial frontal cortex slices; XAβ, custom measurement of immunostained Aβ plaques in fixed medial frontal cortex slices. *We previously generated the Tau and Aβ prion activity measurements in Condello et al. (DOI: 10.1073/pnas.2212954119). ^#^We previously generated the Xtau and XAβ scores in Maxwell et al. (DOI: 10.1186/s40478-021-01298-0).

| Brain bank | Case ID | Case # in study | Age at death | Sex | APOE genotype | Clinical status | Brain region | Postmortem interval (h) | Approx brain mass (g) | Tau prion activity (DxA) | Insoluble total tau (μg/g); formic acid extract<sup>†</sup> | Xtau neuropath score<sup>‡</sup> | Braak stage (NFTs) | Aβ prion activity (DxA) | Insoluble Aβ (μg/g); formic acid extract<sup>‡</sup> | XAβ neuropath score<sup>‡</sup> | CERAD score (amyloid plaques) |
| --- | --- | --- | --- | --- | --- | --- | --- | --- | --- | --- | --- | --- | --- | --- | --- | --- | --- |
| BCN | 907 | 1 | 63 | M | ε3/ε3 | Dementia documented | Frontal cortex | 6 | 1.192 | 253341.25 | 23618 | 4 | VI | 185071.2 | 108 | 3 | C3 |
| UMD | 4870 | 2 | 51 | F | ε2/ε3 | Dementia documented | Frontal cortex | 4 | 1.025 | 179783.64 | 34341 | 4 | ND | 136564.9 | 67 | 4 | ND |
| UMD | 4659 | 3 | 46 | F | ε3/ε4 | No dementia documented | Frontal cortex | 7 | 0.791 | 257720.75 | 102623 | 4 | ND | 170006.7 | 109 | 4 | ND |
| BCN | 714 | 4 | 36 | F | ε3/ε3 | No dementia documented | Frontal cortex | 12 | 1.383 | 172328.76 | 20022 | 2 | II | 100221.7 | 41 | 2 | C2 |

**Supplementary Table S2.** Fixed postmortem donor tissues used for EMBER analysis. Abbreviations: Aβ, amyloid-β; AD, Alzheimer’s disease; APOE, apolipoprotein E; BCN, Barcelona; DS, Down syndrome; EMBER, excitation multiplexed bright emission recording; F, female; KCL, King’s College London; M, male; ND, not determined; OXF, Oxford; UCI, University of California, Irvine; UMAP, uniform manifold approximation and projection; UMD, University of Maryland; Xtau, custom measurement of immunostained neurofibrillary tau tangles in fixed medial frontal cortex slices; XAβ, custom measurement of immunostained Aβ plaques in fixed medial frontal cortex slices. We generated the Xtau and XAβ scores in Maxwell et al. (doi.org/10.1186/s40478-021-01298-0); (−), not determined.

| Brain bank | ID | Disease | Region | UMAP name | Age at death | Sex | APOE | Postmortem interval (h) | Xtau neuropath score | XAβ neuropath score |
| --- | --- | --- | --- | --- | --- | --- | --- | --- | --- | --- |
| KCL | A035/03 | AD | Frontal cortex | AD 1 | 59 | F | ND | 36 | 4 | 4 |
| KCL | A061/03 | AD | Frontal cortex | AD 2 | 55 | M | ND | 18 | 4 | 4 |
| OXF | NP177-2013 | AD | Frontal cortex | AD 3 | 90 | F | ε3/ε3 | 48 | 4 | 3 |
| UCI | 14-08 | AD | Frontal cortex | AD 4 | 86 | M | ε3/ε3 | 4 | 2 | 2 |
| UCI | 37-15 | AD | Frontal cortex | AD 5 | 87 | F | ε3/ε4 | 4 | 2 | 3 |
| UCI | 4-02 | AD | Frontal cortex | AD 6 | 83 | M | ε3/ε4 | 3 | 4 | 3 |
| BCN | 907 | DS | Frontal cortex | DS 1 | 63 | M | ε3/ε3 | 6 | 4 | 3 |
| BCN | 714 | DS | Frontal cortex | DS 2 | 36 | F | ε3/ε3 | 12 | 2 | 2 |
| UMD | 4335 | DS | Frontal cortex | DS 3 | 28 | M | ε4/ε4 | 26 | 2 | 2 |
| UMD | 4870 | DS | Frontal cortex | DS 4 | 51 | F | ε2/ε3 | 4 | 4 | 4 |
| UMD | 5510 | DS | Frontal cortex | DS 5 | 65 | M | ε3/ε3 | 10 | 4 | 4 |
| UMD | 5600 | DS | Frontal cortex | DS 6 | 57 | M | ε3/ε3 | 6 | 4 | 3 |
| UCI | 29-06 | DS | Frontal cortex | - | 45 | F | ε3/ε3 | 3 | 4 | 4 |
| UCI | 29-06 | DS | Temporal cortex | - | 45 | F | ε3/ε3 | 3 | - | - |
| UCI | 3-17 | DS | Frontal cortex | - | 57 | M | ε3/ε3 | 4 | 3 | 4 |
| UCI | 3-17 | DS | Temporal cortex | - | 57 | M | ε3/ε3 | 4 | - | - |
| UCI | 30-00 | DS | Frontal cortex | - | 61 | M | ε3/ε3 | 11 | 4 | 4 |
| UCI | 30-00 | DS | Temporal cortex | - | 61 | M | ε3/ε3 | 11 | - | - |
| UCI | 46-94 | DS | Frontal cortex | - | 62 | F | ε3/ε3 | 3 | 4 | 4 |
| UCI | 46-94 | DS | Temporal cortex | - | 62 | F | ε3/ε3 | 3 | - | - |
| UCI | 30-05 | DS | Frontal cortex | - | 57 | F | ε3/ε3 | 3 | 4 | 4 |
| UCI | 30-05 | DS | Temporal cortex | - | 57 | F | ε3/ε3 | 3 | - | - |

**Supplementary Table S3.** Fresh-frozen postmortem donor tissues used for HTRF and Western blot analysis. Abbreviations: AD, Alzheimer’s disease; ADNC, Alzheimer’s disease neuropathological change; APOE, apolipoprotein E; BCN, Barcelona; DS, Down syndrome; F, female; HTRF, Homogenous Time-Resolved Fluorescence Assay; M, male; MIA, University of Miami; PMI, post-mortem interval in hours; UCI, University of California, Irvine; UMD, University of Maryland.

| Brain bank | Case ID | Cohort | Age | Sex | PMI (h) | APOE | HTRF Tau - Sarkosyl Insoluble |  |  | HTRF Tau - Sarkosyl Soluble |  |  |
| --- | --- | --- | --- | --- | --- | --- | --- | --- | --- | --- | --- | --- |
|  |  |  |  |  |  |  | Replicate 1 | Replicate 2 | Mean | Replicate 1 | Replicate 2 | Mean |
| UMD | 5277 | DS | 19 | M | 26 | 3/4 | 15645.1 | 16458.7 | 16051.9 | 603.1 | 597.5 | 600.3 |
| MIA | 18 | DS | 24 | M | 24 | 3/3 | 545.8 | 542.8 | 544.3 | 542.9 | 542.5 | 542.7 |
| UMD | 5341 | DS | 25 | M | 24 | 3/3 | 553.0 | 548.9 | 550.9 | 545.3 | 545.8 | 545.5 |
| UMD | 4335 | DS | 28 | M | 26 | 4/4 | 2568.1 | 2076.7 | 2322.4 | 554.8 | 559.3 | 557.1 |
| BCN | 714 | DS | 36 | F | 12 | 3/3 | 2166.6 | 5923.8 | 4045.2 | 563.9 | 565.5 | 564.7 |
| UMD | 4904 | DS | 40 | M | 10 | 3/3 | 4267.3 | 4013.1 | 4140.2 | 562.6 | 570.8 | 566.7 |
| UMD | 5783 | DS | 41 | M | 7 | 3/4 | 554.8 | 556.3 | 555.6 | 546.4 | 545.8 | 546.1 |
| UMD | 4659 | DS | 46 | F | 7 | 3/4 | 17361.5 | 15368.8 | 16365.2 | 891.5 | 877.5 | 884.5 |
| UCI | 7-17 | DS | 47 | F | 7 | 3/3 | 7634.3 | 7636.4 | 7635.3 | 2239.4 | 1583.0 | 1911.2 |
| UCI | 32-15 | DS | 49 | M | 6 | 3/3 | 10617.7 | 8329.6 | 9473.7 | 2670.9 | 2536.6 | 2603.7 |
| UMD | 4870 | DS | 51 | F | 4 | 2/3 | 17225.9 | 14728.8 | 15977.4 | 798.9 | 784.0 | 791.4 |
| UCI | 3-17 | DS | 57 | M | 4 | 3/3 | 6796.5 | 7600.6 | 7198.5 | 2059.0 | 1441.0 | 1750.0 |
| UMD | 5600 | DS | 57 | F | 6 | 3/3 | 12901.8 | 15353.2 | 14127.5 | 1162.8 | 1222.3 | 1192.6 |
| UMD | 6151 | DS | 57 | M | 5 | 3/3 | 11733.7 | 8897.7 | 10315.7 | 1796.3 | 1735.5 | 1765.9 |
| UCI | 39-17 | DS | 58 | M | 6 | 3/4 | 13579.0 | 12312.8 | 12945.9 | 1640.6 | 1673.0 | 1656.8 |
| BCN | 1335 | DS | 62 | F | 9 | 3/4 | 5821.2 | 5543.6 | 5682.4 | 1279.2 | 1457.8 | 1368.5 |
| BCN | 1469 | DS | 62 | M | 6 | 3/4 | 16991.4 | 14799.6 | 15895.5 | 709.0 | 699.1 | 704.0 |
| BCN | 907 | DS | 63 | M | 6 | 3/3 | 18942.7 | 17465.2 | 18204.0 | 1065.7 | 1002.4 | 1034.1 |
| UMD | 5386 | DS | 64 | M | 20 | 3/4 | 9503.0 | 8037.1 | 8770.0 | 627.8 | 619.4 | 623.6 |
| UMD | 5510 | DS | 65 | M | 10 | 3/3 | 17315.0 | 14110.2 | 15712.6 | 658.8 | 665.9 | 662.3 |
| UCI | 4-16 | DS | 70 | M | 4 | 3/3 | 6422.3 | 6764.8 | 6593.6 | 1899.6 | 1370.1 | 1634.9 |
| UCI | 21-06 | AD | 82 | M | 5 | 3/4 | 1594.2 | 1561.5 | 1577.9 | 554.0 | 551.8 | 552.9 |
| UCI | 04-02 | AD | 83 | M | 3 | 3/4 | 15727.8 | 17243.0 | 16485.4 | 1100.9 | 1023.3 | 1062.1 |
| UCI | 14-08 | AD | 86 | M | 4 | 3/3 | 17272.3 | 17205.9 | 17239.1 | 719.3 | 685.2 | 702.3 |
| UCI | 37-15 | AD | 87 | F | 4 | 3/4 | 14572.5 | 16351.5 | 15462.0 | 850.5 | 828.6 | 839.6 |
| UCI | 37-15 | AD | 87 | F | 4 | 3/4 | 19684.5 | 20442.3 | 20063.4 | 776.5 | 695.3 | 735.9 |
| UCI | 10-17 | ADNC | 66 | F | 4 | 3/4 | 3615.8 | 2116.9 | 2866.4 | 598.9 | 594.3 | 596.6 |
| UCI | 46-16 | ADNC | 78 | M | 3 | 3/4 | 2028.3 | 1590.5 | 1809.4 | 562.6 | 555.8 | 559.2 |
| BCN | 1937 | ADNC | 83 | F | 8 | 3/3 | 1921.2 | 1603.3 | 1762.2 | 550.5 | 553.6 | 552.1 |
| BCN | 1858 | ADNC | 83 | F | 8 | 3/3 | 14397.1 | 8150.2 | 11273.7 | 570.4 | 604.3 | 587.4 |
| UCI | 18-08 | ADNC | 84 | F | 4 | 3/3 | 606.2 | 619.9 | 613.0 | 548.2 | 549.8 | 549.0 |
| BCN | 1949 | ADNC | 86 | M | 8 | 3/3 | 601.7 | 586.4 | 594.0 | 539.7 | 545.7 | 542.7 |
| UCI | 14-17 | ADNC | 89 | F | 6 | 2/3 | 4252.2 | 4536.0 | 4394.1 | 585.4 | 588.7 | 587.0 |
| BCN | 1870 | ADNC | 97 | F | 7 | 3/3 | 749.4 | 774.4 | 761.9 | 545.5 | 550.4 | 547.9 |

**Supplementary Table S4.** Cryo–electron microscopy data collection and structure determination. Abbreviations: CTE, chronic traumatic encephalopathy; N/A, not applicable; PHF, paired helical filament; SF, straight filament.

|  | Case 1 |  | Case 2 |  | Case 3 |  | Case 4 |  |
| --- | --- | --- | --- | --- | --- | --- | --- | --- |
| Data collection | PHFs | SFs | PHFs | SFs | PHFs | SFs | PHFs | CTE |
| Magnification | x105,000 | x105,000 | x105,000 | x105,000 | x105,000 | x105,000 | x105,000 | x105,000 |
| Defocus range (mm) | -0.8 to -1.8 | -0.8 to -1.8 | -0.8 to -1.8 | -0.8 to -1.8 | -0.8 to -1.8 | -0.8 to -1.8 | -0.8 to -1.8 | -0.8 to -1.8 |
| Voltage (kV) | 300 | 300 | 300 | 300 | 300 | 300 | 300 | 300 |
| Microscope | Titan Krios | Titan Krios | Titan Krios | Titan Krios | Titan Krios | Titan Krios | Titan Krios | Titan Krios |
| Detector | Gatan K3 | Gatan K3 | Gatan K3 | Gatan K3 | Gatan K3 | Gatan K3 | Gatan K3 | Gatan K3 |
| Frame exposure time (s) | 2.024 | 2.024 | 2.024 | 2.024 | 2.024 | 2.024 | 2.024 | 2.024 |
| Dose rate (e-/physical pixel/sec) | 16 | 16 | 16 | 16 | 16 | 16 | 16 | 16 |
| Total dose (e-/Å<sup>2</sup>) | 46 | 46 | 46 | 46 | 46 | 46 | 46 | 46 |
| Pixel size (Å) | 0.834 | 0.834 | 0.834 | 0.834 | 0.834 | 0.834 | 0.834 | 0.834 |
| Movies collected | 9160 |  | 5902 |  | 6949 |  | 1293 |  |
| Grids & sample | Conventional grids<br>Pronase |  | Affinity grid<br>no Pronase |  | Conventional grid<br>no Pronase |  | Affinity grid<br>no Pronase |  |
| Reconstruction |  |  |  |  |  |  |  |  |
| Box size (pixel) | 280 | 280 | 280 | 280 | 280 | 280 | 280 | 280 |
| Inter-box distance (Å) | 17 | 17 | 17 | 17 | 17 | 17 | 17 | 17 |
| Total segments* | 1127462 | 329537 | 7488898 | 287943 | 271767 | 86388 | 197928 | 17473 |
| Final Particles (no.) | 215640 | 47801 | 79599 | 26964 | 47264 | 35177 | 32831 | N/A** |
| Resolution (Å) | 2.7 | 3 | 3.1 | 3.2 | 2.9 | 3.1 | 5 | 7.8 |
| B-factor (Å<sup>2</sup>) | -83.71 | -81.34 | -89.69 | -71.29 | -68.79 | -90.05 | -262.95 | -415.37 |
| Helical rise (Å) | 2.39 | 4.81 | 2.37 | 4.77 | 2.4 | 4.81 | 2.38 | 2.38 |
| Helical twist (°) | 179.45 | -1.08 | 179.5 | -1.04 | 179.48 | -1.06 | 179.45 | 179.55 |

\*after initial 2D classification
\*\*no subsequent classification

**Supplementary Table S5.** Cryo–electron microscopy model building. Abbreviations: #, number; PHF, paired helical filament; RMSD, root mean square deviation; SF, straight filament.

|  | Case 1 |  | Case 2 |  |
| --- | --- | --- | --- | --- |
| Model | PHFs | SFs | PHFs | SFs |
| model resolution | 2.7 | 3.0 | 3.1 | 3.5 |
| atoms | 35910 | 39501 | 40698 | 39513 |
| Residues | 2310 | 2541 | 2618 | 2541 |
| Bonds (RMSD) |  |  |  |  |
| Length (Å) (# > 4σ) | 0.010 (0) | 0.010 (0) | 0.010 (0) | 0.011 (0) |
| angles (°) (# > 4σ) | 1.717 (92) | 1.791 (70) | 1.871 (38) | 1.914 (95) |
| MolProbity score | 2.21 | 2.32 | 2.53 | 2.63 |
| Clash score | 18.87 | 32.66 | 40.97 | 46.84 |
| Ramachandran plot (%) |  |  |  |  |
| Outliers | 0.00 | 0.00 | 0.00 | 1.33 |
| Allowed | 6.67 | 4.69 | 6.67 | 6.67 |
| Favored | 93.33 | 95.31 | 93.33 | 92 |
| Rotamer outliers (%) | 0.00 | 0.00 | 0.00 | 0.00 |

